# Automating Glycosylation of Protein Models in the Martini 3 Force Field with GlycoMarinate

**DOI:** 10.64898/2026.11.20.768051

**Authors:** Alam Ahmad Hidayat, Alexis Descamps, Krzysztof Murzyn, Balázs Fábián, Mateusz Sikora

## Abstract

Most membrane and secreted proteins undergo glycosylation, the covalent posttranslational addition of oligosaccharide chains. Glycans, which constitute up to 30% of a glycoprotein’s mass, alter protein trafficking, dynamics, interactions, and thus function. Because of their heterogeneity, rapid dynamics, and conformational richness, glycans are difficult to characterise experimentally, so molecular simulations can provide a powerful alternative for studying their effects on proteins. Given that the time and length scales involved are typically large, coarse-grained models can be used to enhance the sampling and allow for the study of interactions of glycoproteins in their cellular context. Although individual saccharides and glycans have been parameterised for coarse-grained simulations, modelling new glycan structures has largely required system-specific parameter development, and no general framework connects transferable glycan models with automated construction of complete glycoproteins. Here, we introduce GlycoMarinate, a general framework for coarse-grained modelling of N-glycoproteins within the Martini 3 force field. The framework supports two complementary levels of glycan description: a transferable generic model designed for broad applicability, and refined glycan-specific models optimised against atomistic conformational ensembles. We derive and validate refined models for ten representative high-mannose, hybrid, and complex N-glycans. Both descriptions are integrated into an automated workflow that generates the topologies and clash-free starting coordinates for site-specifically glycosylated proteins. We demonstrate the framework on biologically relevant systems, including the densely glycosylated SARS-CoV-2 spike protein.

## 1 Introduction

Most proteins undergo post-translational modifications (PTMs), which expand molecular diversity beyond that encoded by the genome and confer tissue-specific functional properties. Among them, glycosylation—the covalent addition of complex carbohydrates to selected amino-acid side chains in proteins—is the most ubiquitous and diverse modification, and is also often the largest by mass of all known PTMs. Glycans, which decorate the majority of membrane and secreted proteins [1], can be divided into two broad classes with distinct biological and chemical properties. O-linked glycans are attached to proteins through an O-glycosidic bond formed with the hydroxyl group of serine (Ser) or threonine (Thr) residues. This group contains also the only known cytoplasmic form of glycosylation — O-GlcNAcylation — involving the reversible addition of a single N-acetylglucosamine (GlcNAc) residue to cytoplasmic and nuclear proteins. N-linked glycans are typically attached through an N-glycosidic bond to the amide nitrogen of asparagine (Asn) within the Asn–X–Ser/Thr consensus sequence, where X is any amino acid except proline.

N-linked glycans share a conserved core consisting of two GlcNAc and three mannose (Man) residues, which can be elaborated into high-mannose, hybrid, or complex architectures. These structures differ in both monosaccharide composition and topology: high-mannose glycans remain enriched in Man residues, hybrid glycans combine a mannose-rich branch with a branch undergoing complex-glycan maturation, whereas fully mature complex glycans contain bi-, tri-, or tetra-antennary branches extended with GlcNAc, galactose (Gal), fucose (Fuc), and sialic acid residues, predominantly N-acetylneuraminic acid (Neu5Ac). Consequently, N-glycan identity is defined not only by composition but also by its branching pattern, linkage position, anomeric configuration, and terminal capping patterns, which might differ widely across organisms, cell types and disease states.

Glycans are assembled and processed mainly in the endoplasmic reticulum (ER) and Golgi apparatus, where they contribute to protein folding, quality control, maturation, and trafficking. On mature proteins, glycans modulate conformational dynamics, solubility, aggregation, and molecular recognition, shaping protein–protein, protein–membrane, and host–pathogen interactions. They are central to self-recognition and immune regulation, while pathogens such as enveloped viruses exploit host-derived glycans to mask antigenic surfaces, as exemplified by the dense glycan shield of the SARSCoV-2 Spike protein [2]. Aberrant glycosylation is also a hallmark of numerous human diseases and often contributes directly to pathogenesis [3–6].

Yet, despite their broad functional and pathological importance, connecting glycan composition to molecular mechanism remains difficult. Such a connection requires a structural description of the conformational ensembles populated by glycans and of their interactions with the surrounding molecular environment. This is intrinsically challenging because glycans are highly flexible molecules. They populate multiple conformational states separated by relatively shallow energy barriers, enabling rapid interconversion between conformers in aqueous solution [7]. Together with their chemical heterogeneity, these conformational dynamics complicate experimental structure determination, and all-atom molecular dynamics (AA MD) simulations have therefore become a powerful approach to study their effects on protein dynamics and interactions. However, AA MD remains prohibitively expensive for large, multicomponent glycoprotein systems and subcellular environments, where glycan-mediated interactions and collective behaviour become important. Thus, coarse-grained (CG) models, in which groups of atoms are represented by effective interaction sites, offer an attractive strategy to extend simulations toward mesoscopic time and length scales required to capture those effects.

Among CG approaches, Martini [8] has emerged as the leading biomolecular force field. The chemical character of the underlying atomistic groups is encoded by assigning each CG interaction site a Martini bead type, which defines its non-bonded interaction parameters and therefore the Lennard– Jones (LJ) interaction strength with other bead types. Initially developed for lipids [9], Martini has progressively expanded and refined its treatment of lipid membranes [10] evolving into a versatile framework for proteins [8], nucleic acids [11], carbohydrates [12], solvents, and biologically relevantions, thereby enabling simulations of complex biomolecular assemblies. However, glycoproteins remain comparatively difficult to model within the current Martini implementation. In Martini 2, carbohydrate models were developed for several saccharides and manually parametrized glycans [13–15], but their broader adoption was limited by known interaction-balance problems, including excessive effective attraction [16] and between saccharides and proteins [**?**], as well as by stability issues and the absence of a systematic glycoprotein-building workflow.

Martini 3 now provides a substantially improved carbohydrate foundation [12], including transferable mapping and parametrization rules for saccharides derived and validated against atomistic reference models. These advances have enabled systematic efforts to generate disaccharide parameters [17] and inspired more complex linear polysaccharide models [18,19]. More recently, the disaccharide-based approach has been implemented for building CG parameters for multiple lipopolysaccharides found in common pathogens [20]. However, they do not yet provide a practical route to simulation-ready glycoproteins, which requires integrating glycan parametrization, protein linkage, and generation of clash-free starting structures. Consequently, glycosylation remains difficult to incorporate routinely into Martini simulations.

Here, we introduce GlycoMarinate, a general framework for Martini 3 simulations of Asn-linked N-glycoproteins that integrates transferable glycan parametrization with automated construction of simulation-ready glycoprotein models. At its core, the framework provides a generic parametrization strategy based on a compact set of transferable local bonded terms (GlycoMarinate Transferable model, **GMT**), enabling a broad range of common non-bisecting N-glycan architectures to be constructed without glycan-specific parametrization. For applications requiring a more detailed reproduction of glycan conformational ensembles, we additionally derive optimized, glycan-specific model (GlycoMarinate Refined model, **GMR**) for ten representative high-mannose, hybrid, and complex N-glycans.These force-field descriptions are coupled to a modular, open-source construction workflow that generates the GlcNAc–Asn linkage, glycoprotein topology, and clash-free starting coordinates for user-defined sitespecific glycosylation patterns. Finally, we validate the generic and refined glycan models against atomistic simulations and available experimental data and demonstrate the complete framework on biologically relevant glycoproteins, including the densely glycosylated SARS-CoV-2 Spike protein.

## 2 Results

### 2.1 GlycoMarinate integrates N-glycan models with automated glycoprotein construction

GlycoMarinate is organized into two coupled layers: glycan force-field description and glycoprotein construction. The first supports the GMT model assembled from transferable interaction terms and the GMR model, with glycan-specific bonded terms optimized against atomistic conformational ensembles for a representative set of glycans (Figure 1A). The second layer transforms site-specific glycosylation assignments into simulation-ready glycoprotein models by generating the GlcNAc–Asn connectivity, complete topology, and sterically viable starting coordinates (Figure 1B, C). We first describe how these force-field descriptions are deployed to construct complete glycoprotein models, before examining their parametrization and validation in the following sections.

**Figure 1:**
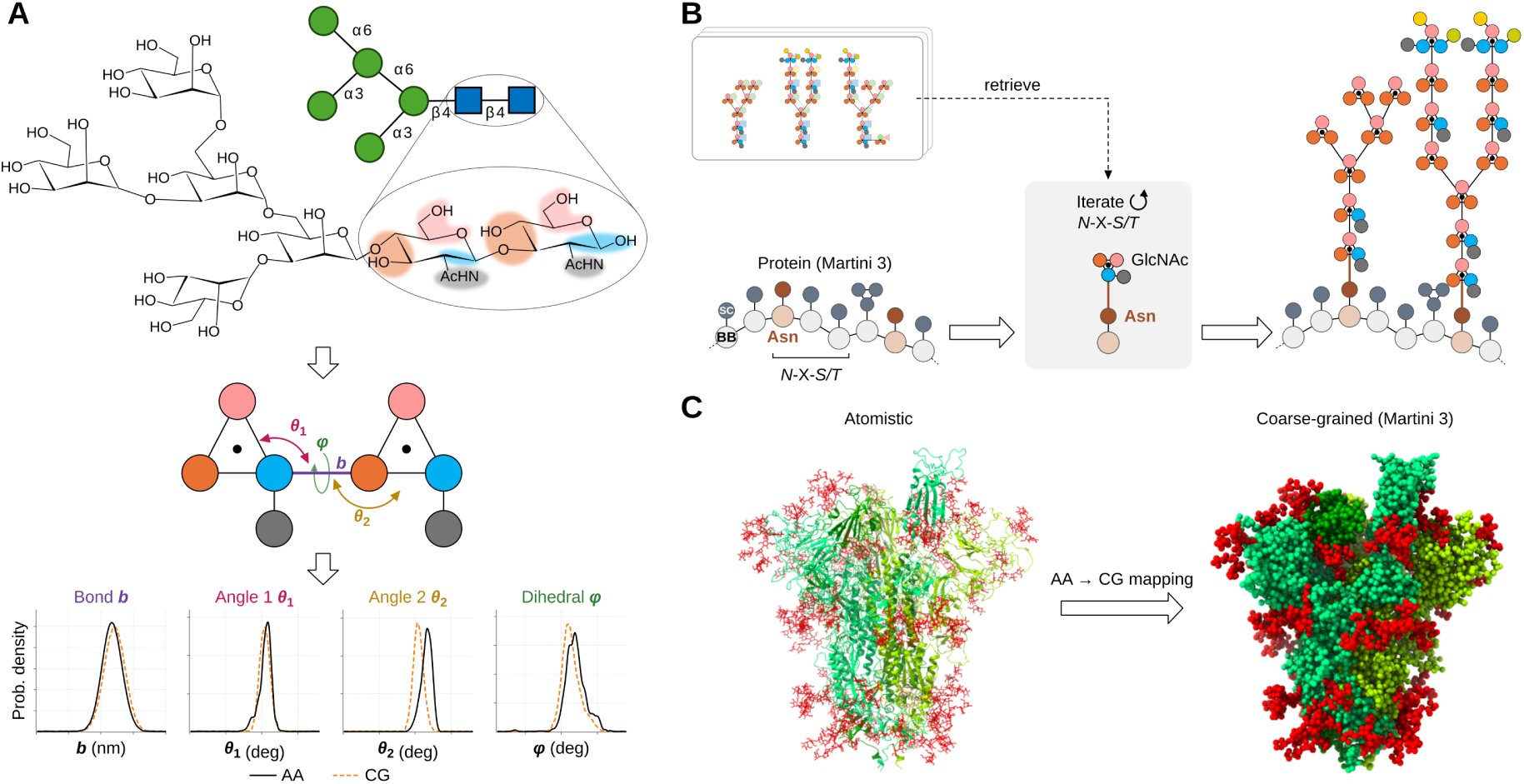
Overview of the GlycoMarinate framework for Martini 3 N-glycoprotein modelling. **A.** Glycan parametrization. Representative high-mannose, hybrid, and complex N-glycans are mapped to Martini 3 beads. Glycosidic linkages are described by local bonded terms derived from mapped glycan disaccharide fragments, supplemented by angular terms spanning three consecutive monosaccharides. These transferable local terms form the basis of the GMT model, whereas glycan-specific bonded parameters in the GMR model are optimized against all-atom reference ensembles mapped to the coarse-grained representation. **B.** Topology construction. An existing Martini 3 protein topology is combined with user-defined N-glycosylation sites and the corresponding glycan topologies through the parametrized GlcNAc–Asn linkage to generate a complete glycoprotein topology. **C.** Coordinate construction. An atomistic protein structure can be mapped to Martini 3, or an existing CG protein model can be used directly. Compatible atomistic glycan conformers are mapped and combined with the protein coordinates, with optional generation of clash-free starting configurations, to yield a simulation-ready N-glycoprotein model.

The glycan parametrization layer comprises two complementary levels. The GMT model builds N-glycans from transferable bonded terms focused around inter-saccharide linkages, extending the fragment-based Martini 3 carbohydrate representation [12] with disaccharide-level parameters (Figure 1A). In contrast, the GMR mode uses glycan-specific bonded parameters optimized against mapped atomistic conformational ensembles. We derived such models for representative high-mannose, hybrid, and complex N-glycans. Thus, GlycoMarinate can either assemble diverse glycans from reusable local terms or use optimized parameters.

Constructing a Martini 3 representation of an N-glycoprotein requires coordinated treatment of three elements: a protein topology, one or more glycan topologies, and parameters for a covalent GlcNAc-Asn linkage to the protein. GlycoMarinate automates this integration by combining the protein topology with the assigned glycan topologies and introducing the appropriate GlcNAc–Asn connectivity, at each user-defined glycosylation site. The procedure starts from an existing Martini 3 protein topology, which can be generated using *martinize2*. GlycoMarinate extends this topology by adding the glycan topologies assigned to user-defined N-glycosylation sites and introducing the predefined GlcNAc–Asn connectivity. Because glycosylation is introduced as an extension of the existing protein topology, protein-specific refinements are preserved, including, for example, Go-Martini [21] or modified Martini 3 models for intrinsically disordered proteins [22]. Glycan topologies are added iteratively via *polyply* [23]. Although the current implementation provides GMT and GMR models, the topology construction procedure can accommodate compatible user-provided glycan models.

For coordinate generation, the workflow is agnostic to the source of the initial glycan conformations and requires only a sterically viable atomistic glycoprotein structure consistent with the specified glycosylation pattern. Such coordinates can originate from an existing model or from external glycoprotein-building and conformer-generation tools [24,25]. In the present workflow, we adopt GlycoSHIELD [26] as a convenient default to graft glycan conformers onto the protein while filtering protein–glycan and glycan–glycan clashes. The provided glycan structures are then mapped to Martini resolution using the predefined atom-to-bead mapping and combined with the corresponding coarse-grained protein coordinates to produce the complete starting structure. In the following sections, we define and validate the glycan mapping, GlcNAc–Asn linkage, and bonded interactions underlying the generic and refined models.

### 2.2 A representative N-glycan panel in GlycoMarinate

In this work, we employed a compact panel of ten non-bisecting N-glycans spanning the major structural features relevant to Martini 3 parametrization (Figure 2**D**). The set includes high-mannose, hybrid, and complex architectures and samples core fucosylation, terminal sialylation, and increasing antenna number. Together, these structures span repeated mannose linkages, chemically distinct antennae, charged terminal residues, and progressively more complex branching patterns. They therefore provide the linkage, branching, and size diversity needed to establish the common mapping and linkage rules, optimize glycan-specific bonded interactions, and assess the transferability of the generic fragment-based model. Bisecting GlcNAc-containing structures were not included in the present parametrization and therefore remain outside the validated scope of the framework. Despite relative richness of the glycan types and shapes, the striking repetitiveness of the linkage types suggests that a transferable, fragment-based approach could be used to parametrize these glycans in Martini.

**Figure 2:**
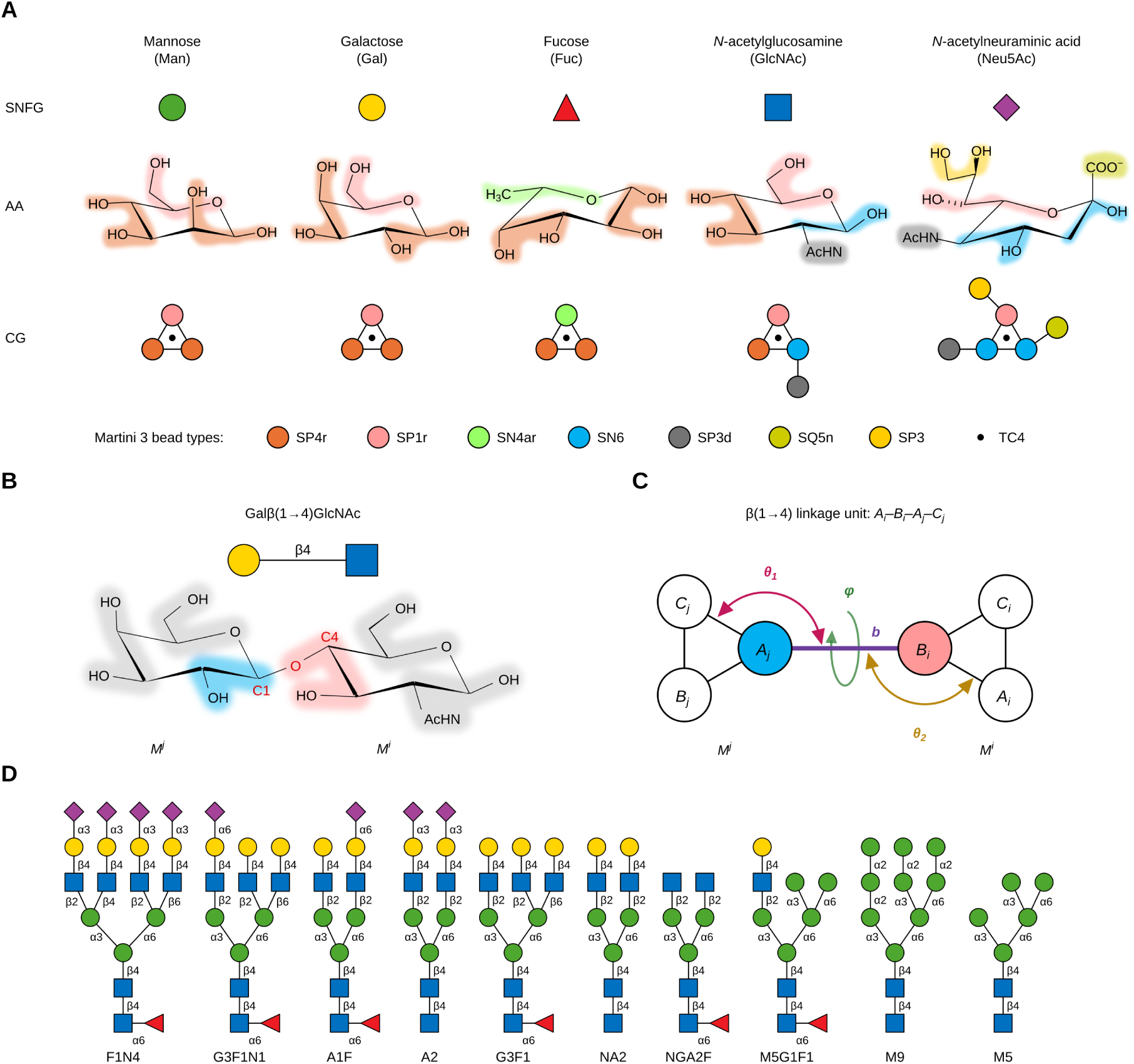
Linkage-aware mapping and bonded representation of N-glycans in GlycoMarinate. **A.** Monosaccharide representation across levels of resolution. Representative residues are shown using Symbol Nomenclature for Glycans (SNFG), followed by their atomistic structures with the Martini 3 atom-to-bead mapping overlaid. **B.** Glycosidic bond formation modifies the composition, polarity, and hydrogen-bonding capacity of the mapped fragments participating in the linkage. **C.** Coarse-grained representation of the linkage unit and with its local bond, angle, and dihedral terms **D.** Panel of ten non-bisecting N-glycans used in this work, spanning representative high-mannose, hybrid and complex architectures.

### 2.3 N-glycan parametrization strategy in Martini 3 force field

#### Linkage-and branch-aware mapping enables modular Martini 3 N-glycan construction

We next propose a mapping scheme for N-glycans that preserves compatibility with Martini 3 carbohydrate conventions while enabling automated construction of branched glycan topologies. By mapping, we understand representing a group of 2-5 heavy atoms as a CG bead placed at the centre of geometry (CoG) of such group. As defined by Grünewald et al. [12], transferable mapping for carbohydrates requires adhering to three rules: 1) maximize the number of canonical four-heavy-atom mappings (i.e., four heavy atoms per CG bead); 2) avoid splitting functional groups across multiple beads; 3) start mapping at the anomeric carbon and proceed in ascending carbon number until all atoms are assigned to beads.

Following this notion, pyranose-like rings are therefore represented by three small (S) or tiny (T) beads arranged into a triangle, with an additional virtual site placed at the triangle’s CoG (TC4 bead type). Such a geometry was found to preserve the saccharide ring planarity best and facilitate stacking with aromatic groups [27]. Such choice also improves the matching of non-hydrogen atoms to beads — the recommended in martini 3 ratio of at most one unmatched hydrogen per ten non-hydrogen atoms mapped to CG beads [21]. The CG constructions of five monosaccharides represented in the selected panel of N-glycans are shown in Figure 2**A**.

Direct reuse of isolated monosaccharide mappings is insufficient for polysaccharides because glycosidic bond formation removes a hydroxyl group from the donor anomeric carbon and a hydrogen atom from the acceptor hydroxyl group. To address this, we define each glycosidic “linkage unit” as a pair of linkage-aware bead fragments, written as *M^j^a*(*J → I*)*M^i^*, where *M^i^* and *M^j^* are two connected monosaccharides indexed *i* and *j*, with *j > i* (residues numbered from the reducing end). Here, *a* denotes the *α* or *β* anomeric configuration, while *I* and *J* denote the carbon locants (International Union of Pure and Applied Chemistry — IUPAC — carbohydrate numbering) of the acceptor and donor residues, respectively. *J* = 1 for all donors except Neu5Ac, for which *J* = 2. Atom assignment follows the CHARMM-compatible convention for the (C*J*–O–C*I*) linkage pattern, in which the glycosidic oxygen belongs to the acceptor residue *M^i^*. For example, in the Gal*β*(1 *→* 4)GlcNAc linkage unit (see Figure 2B), the anomeric carbon (C1) belongs to galactose (monomer *M^j^*), while the glycosidic oxygen is part of GlcNAc (monomer *M^i^*). Beads within each monosaccharide *M^k^* are labelled alphabetically (*A*, *B*, *C*, etc.) following the locant order of the atom-to-bead mapping. Bead *X* of monosaccharide *M^k^* is denoted *X_k_*; for example, *A_i_* denotes bead *A* of monosaccharide *M^i^*. This scheme standardizes the mapping across linkage types and avoids mixing atoms from neighbouring residues.

Glycosidic bond formation removes a hydroxyl group or a hydroxyl hydrogen from the beads containing the linking atoms, reducing their polarity and hydrogen-bonding capacity; these beads are therefore reassigned relative to the isolated monosaccharides (Supplementary Table S1). In the donor *M^j^*, the anomeric bead *A_j_* loses O1 and retains C1, C2 and O2, and its SP4r type is replaced by SN6. In the acceptor *M^i^*, the bead containing the glycosidic oxygen loses only the hydroxyl hydrogen and is assigned a less polar type: SP4r becomes SP1r for *I* = 3 or 4 (bead *B_i_*), and SP1r becomes SN6r for *I* = 6 (bead *C_i_*). For *I* = 2, the acceptor bead *A_i_* is also the anomeric bead of *M^i^* and is treated as described below for branching mannoses. Two donor residues deviate from this scheme. In GlcNAc, C2 carries the N-acetyl group, which is mapped to a separate bead, so the anomeric bead of the isolated monosaccharide contains only C1, C2 and O1, an ethanol-like fragment. Removal of O1 leaves an alkane-like two-carbon fragment, which is represented by a TC3 bead, the default for analogous cyclic alkane-like fragments in Martini 3 small-molecule parametrizations [28]. In Neu5Ac, whose anomeric carbon is C2, the anomeric bead comprises the carboxylate group (C1, O11, O12) together with C2 and C3 and is assigned a regular-size charged Q5n bead [12]. The same strategy extends naturally to branching points. In non-bisecting N-glycans, the conserved core is a pentasaccharide composed of two GlcNAc and three Man residues. Within this core, the *β*-mannose acts as the primary branching point, carrying two arm mannoses through *α*(1 *→* 3) and *α*(1 *→* 6) linkages.

The arm mannoses can be elongated by additional monosaccharides; for example, each arm of the A2 glycan carries a Neu5Ac*α*(2 *→* 3)Gal*β*(1 *→* 4)GlcNAc*β*(1 *→* 2) sequence. In triand tetra-antennary glycans, one or both arm mannoses additionally act as secondary branching points. When a mannose acts as a donor through C1 and as an acceptor at O2, both linkage sites fall in bead *A*, which requires an adjusted mapping. For example, bead *A* of the *α*(1 *→* 6) arm mannose loses not only the O1 hydroxyl group upon formation of the Man*α*(1 *→* 6)Man bond but also the O2 hydroxyl hydrogen upon formation of the GlcNAc*β*(1 *→* 2)Man bond. This bead therefore contains only C1, C2, and O2, and is assigned SN5 instead of SN6 to reflect its further reduced polarity (Supplementary Figure S1). The same scheme applies to the mannose in the *α*(1 *→* 3) arm.

We encoded the resulting mapping rules as an atom-to-bead dictionary covering the monosaccharides, linkage types, and branch-specific substitutions in the selected N-glycan panel (Supplementary Table S1). GlycoMarinate uses this dictionary internally to map atomistic glycan coordinates to coarsegrained resolution. We retain the CHARMM atom-naming convention for consistency with previous Martini 3 carbohydrate models [12], the CHARMM-GUI Martini Maker [29], and the CHARMM36 carbohydrate force field [30, 31] used to generate the GlycoSHIELD reference conformers [26]. The pipeline itself is, however, agnostic to atom naming; switching to another convention, such as that of GLYCAM [32], requires only renaming the atoms in the mapping file.

### Selecting stable bonded-term configurations for glycosidic linkages

We parametrized only the bonded terms associated with glycosidic linkages and retained the intramonomer terms of the Martini 3 monosaccharide models [12], including ring-bead constraints and the angles and dihedrals of functional groups. Each linkage unit is described by four bonded terms (Figure 2**C**): one bond, two angles, and one dihedral spanning the same four beads. Previous Martini 3 carbohydrate models used four angles and two dihedrals per linkage [12, 17]; in branched N-glycans, we found that this scheme leads to numerical instabilities at branching points (Supplementary Figure S2**A**). Because several linkage units share beads at these points, their dihedrals impose overlapping torsional restraints on the same beads, generating competing forces that destabilize the simulations. To identify a robust configuration for each linkage unit, we screened candidate choices of the four beads defining the bonded terms. M9 and F1N4 were used as test cases because both contain primary and secondary branching points. Candidate configurations were first assessed in 1 *µ*s simulations of these two glycans, and those yielding stable trajectories were then tested on the remaining glycans. Screening stopped at the first configuration that remained stable for all ten glycans, as demonstrated for *α*(1 *−* 6) linkage determination in Supplementary Figure S2**B**. The similar search strategy was adopted for inferring dihedral configurations for other linkage types. The resulting bead configurations depend only on the linkage type, defined by the anomeric configuration and the acceptor locant, and not on the identity of the monosaccharides (Table 1).

**Table 1:** Bead configurations defining the bonded interactions of glycosidic linkages at the CG level. Each row lists four beads (1–2–3–4), with the acceptor *M^i^* on the left and the donor *M^j^* on the right. The solid line marks the glycosidic bond (2–3), and the dashed lines complete the two linkage angles (1–2–3 and 2–3–4); the dihedral spans all four beads. *X_k_* denotes bead *X* of monosaccharide *M^k^*. Neu5Ac*α*(2 → 3) and Neu5Ac*α*(2 → 6) linkages use the *α*(1 → 3) and *α*(1 → 6) configurations, respectively.

| Linkage | *M*<sup>*i*</sup> |  |  | —— | *M*<sup>*j*</sup> |  |  |
|---|---|---|---|---|---|---|---|
|  | 1 | ---- | 2 | —— | 3 | ---- | 4 |
| *α*(1 → 2) | *B*<sub>*i*</sub> | ---- | *A*<sub>*i*</sub> | —— | *A*<sub>*j*</sub> | ---- | *C*<sub>*j*</sub> |
| *α*(1 → 3) | *A*<sub>*i*</sub> | ---- | *B*<sub>*i*</sub> | —— | *A*<sub>*j*</sub> | ---- | *B*<sub>*j*</sub> |
| *α*(1 → 6) | *A*<sub>*i*</sub> | ---- | *C*<sub>*i*</sub> | —— | *A*<sub>*j*</sub> | ---- | *B*<sub>*j*</sub> |
| *β*(1 → 2) | *C*<sub>*i*</sub> | ---- | *A*<sub>*i*</sub> | —— | *A*<sub>*j*</sub> | ---- | *C*<sub>*j*</sub> |
| *β*(1 → 4) | *A*<sub>*i*</sub> | ---- | *B*<sub>*i*</sub> | —— | *A*<sub>*j*</sub> | ---- | *C*<sub>*j*</sub> |
| *β*(1 → 6) | *A*<sub>*i*</sub> | ---- | *C*<sub>*i*</sub> | —— | *A*<sub>*j*</sub> | ---- | *C*<sub>*j*</sub> |

### Generic bonded parameters of N-glycans in Martini 3

For the GMT parametrization strategy, we first enumerated all unique linkage units across all glycans in Figure 2**D**. For each unit, we identified all glycans containing the linkage, defined four geometric variables based on the dihedral fingerprint lookup (Table 1), and obtained their distributions of AA trajectories mapped to CG resolution (pseudoCG trajectories). Next, the distributions for each variable were pooled with equal weight across all glycans. Using the Boltzmann inversion [33], the resulting distributions were fitted to the appropriate potential functions (Equations 1-4). For bimodal bond and angle distributions, we fitted only to the highest and narrowest mode. Moreover, the linkage angles with distribution peaks located beyond 135° were automatically fitted with the restricted bending potential (Equation 3) to avoid instability from near-collinear dihedral configurations. The resulting fitted parameters were adopted as the generic parameters for that particular linkage unit. We iterated the procedure for 12 unique linkage units, as illustrated in Figure 3**A**. The final generic linkage parameters are reported in Table S2. To construct the CG topology of a given N-glycan, a monomer-level graph was manually defined by assigning a sequential index to each monomer and identifying adjacent monosaccharide pairs and their linkage types. From this, each linkage unit was matched to the generic linkage parameter library from the previous stage. The parameters for each unit were retrieved from the library and written as a topology output. The final CG glycan topology was finalized by adding intramonomer parameters from individual monosaccharides (Figure 2**A**), including ring distance constraints and bonded parameters for internal functional groups. This modular strategy can be implemented for building any N-glycan parameters with different sizes and complexity. In Figure 3**B**, we illustrated this for A1F glycan. By retrieving 8 unique linkage units, the topology was stitched sequentially with the appropriate linkage parameters based on its monomer-level connectivity. The order of the monosaccharide pairs can be inferred manually from its SNFG representation or supplemented from its condensed IUPAC notation.

**Figure 3:**
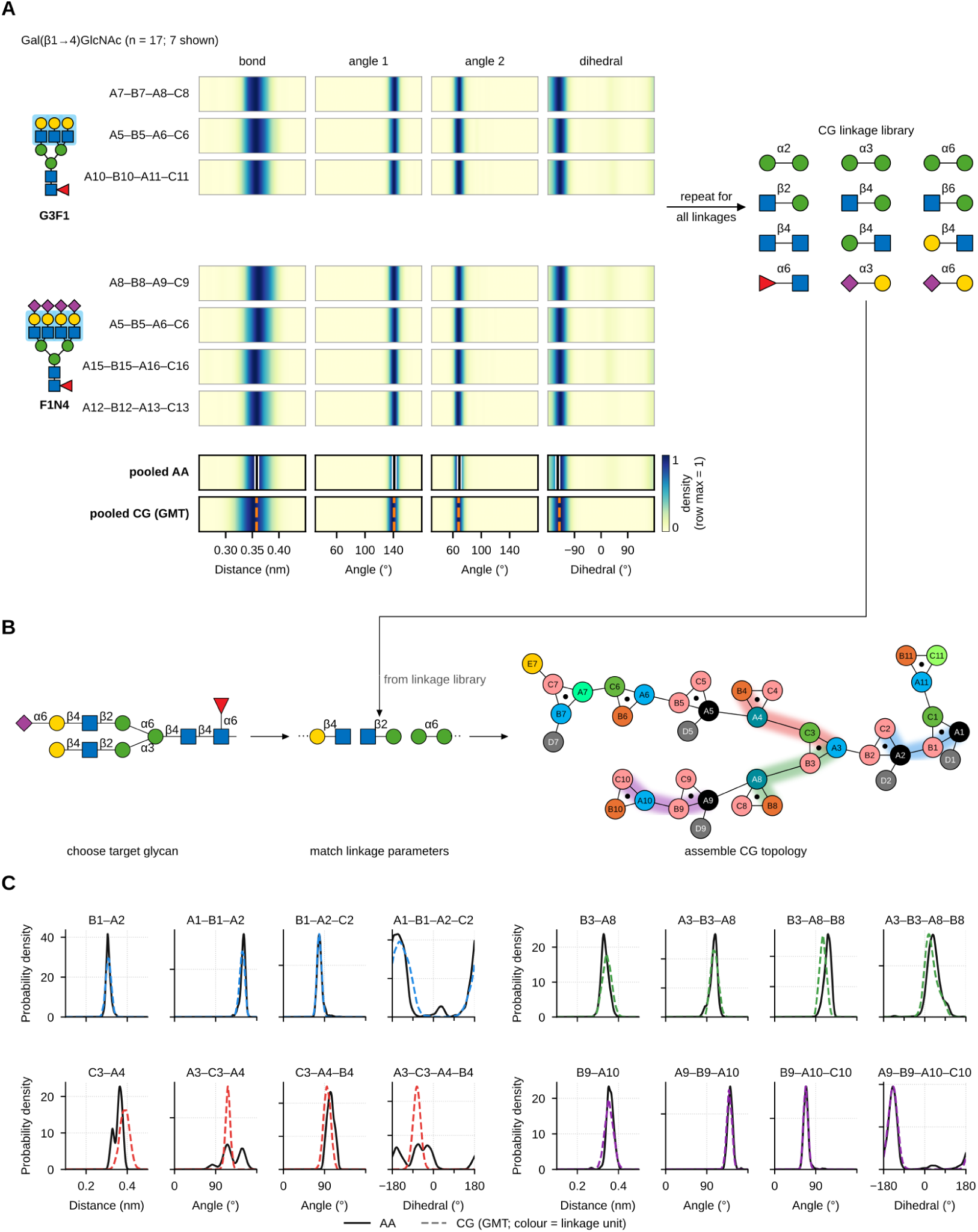
Generic parametrization of glycosidic linkages and assembly of glycan topologies. **A.** Linkage parametrization, shown for Gal*β*(1 4)GlcNAc. Each row shows the mapped AA distributions of the bond, the two angles and the dihedral of one instance of the linkage unit, labelled by its bead quadruple (e.g. A7–B7–A8–C8); 7 of the 17 instances in the glycan panel are shown, from G3F1 and F1N4 (blue shading in the SNFG sketches). Each strip is normalised to its maximum (colour bar). The pooled AA row combines all 17 instances with equal weight (black line, AA mean), and the pooled CG row shows the CG simulations with the generic (GMT) parameters (dashed orange line, minimum of the generic potential). Repeating this for every linkage type gives the CG linkage library of 12 units. **B.** Glycan parametrization, shown for A1F: the linkage units of the target glycan are matched to the library and combined with the intramonomer terms into the CG topology. Beads are coloured by type as in Fig. 2A; coloured shading marks the four linkage units shown in C. **C.** Parameter validation for A1F: distributions of the bond, two angles and dihedral of the four linkage units highlighted in B, from the AA reference (solid black) and the GMT model (dashed, coloured as the corresponding unit in B).

Using this topology, we performed 3 *µ*s CG simulations for all glycans to extract their probability parameter distributions. The resulting CG distribution plots were compared against AA CHARMM-36 distributions, as shown in Figures S3-S12. Across all glycans, the CG distributions showed good visual agreement with the AA references, with closely aligned peaks in bond and angle distributions. For A1F, which comprises eleven monomers and eight unique linkage units, the CG distributions agreed well with the AA reference across all linkage units (Fig. 3**C**). The main discrepancies occurred for the Man*α*(1 *→* 6)Man linkage, where three of the four terms differ from the AA reference. The AA bond and the *A*_3_–*C*_3_–*A*_4_ angle are multimodal, whereas the CG model samples a single, slightly longer bond and a single angle between the AA modes. The CG dihedral is also much narrower than the broad, multimodal AA distribution. Only the second angle, *C*_3_–*A*_4_–*B*_4_, is well reproduced. The AA distributions of this linkage vary between glycans, depending on whether the arm carries a *β*(1 *→* 6)-linked GlcNAc or a sialylated antenna or sits next to a fucosylated core. A single generic term therefore represents only the majority state.

Moreover, we also computed the mean distance score between the two distributions, proposed by Bresnett et al. [34] and termed in this work as the Fast Forward (FF) score (see Method, eq. 6). Lower FF scores indicate a greater agreement between AA and CG distributions. The resulting FF scores for all glycans are shown in Table 2, with total average scores are within a range of 0.2 to 0.4. Imposing 0.5 as the acceptance threshold, at least 90% of bonded distributions were adequately reproduced by the generic parametrization. Moreover, the breakdown into bonds, angles, and dihedral scorings (Figure S14**B**) reveals that the dihedral agreement is marginally better (all scores are *<* 0.4) than the bond and angle distributions; with 7 glycans even scored below 0.3, indicating excellent agreement. We observed that the discrepancy in bonds and angles likely arose from the generally sharper unimodal distributions in the pseudo-CG (AA) reference compared with the CG simulations, where small maxima deviations could be penalized more strongly, yielding larger FF scores.

**Table 2:** Total FF score (mean SD over all bonded parameters) of the generic transferable (GMT) and refined (GMR) CG models relative to the atomistic reference contributed by *n* bonded variables per glycan. Lower values indicate better agreement.

| Glycan | *n* | GMT | GMR |
|---|---|---|---|
| M5 | 24 | 0.34 ± 0.21 | 0.25 ± 0.10 |
| NGA2F | 28 | 0.24 ± 0.12 | 0.21 ± 0.07 |
| M5F1G1 | 36 | 0.28 ± 0.16 | 0.24 ± 0.13 |
| NA2 | 32 | 0.23 ± 0.12 | 0.21 ± 0.08 |
| M9 | 40 | 0.33 ± 0.21 | 0.23 ± 0.09 |
| A1F | 40 | 0.22 ± 0.10 | 0.22 ± 0.10 |
| G3F1 | 44 | 0.23 ± 0.12 | 0.23 ± 0.09 |
| G3F1N1 | 48 | 0.26 ± 0.16 | 0.21 ± 0.09 |
| A2 | 40 | 0.31 ± 0.20 | 0.28 ± 0.11 |
| F1N4 | 68 | 0.26 ± 0.17 | 0.22 ± 0.10 |

Especially for bond length in the M5 glycan, the observed deviation was most drastic (FF score = 0.51 in Figure S14). This largely originates from the differences in mapping Man(*α*1 *−* 3)Man and Man(*α*1 *−* 6)Man in branch regions between high-mannose glycans (M5, M9, and M5F1G1) and the complex glycans (Figure S17). In complex glycans, the primary branching region involves *β*-linked Man-GlcNAc units, whereas the corresponding region in high-mannose glycans involves *α*-linked Man–Man units. Differences in hydrogen and hydroxyl group arrangements shift the mapped center of geometry (CoG), resulting in a slightly misaligned distribution peak of CG bond length between the two glycan types. Because our panel is dominated by complex glycans (7 glycans), the maxima of bond length were biased towards those complex glycans. Consequently, applying these parameters to M5 resulted in a slight discrepancy shown in FF scoring for the glycan.

Nevertheless, the average bond and angle scores across glycans remained below 0.5 (acceptable), supporting the visual agreement of the distributions shown in Figures S3-S12. The overall agreement supports the generic parametrization as a reasonable first approximation for N-glycans. Additionally, we also reported the normalized Wasserstein distance (nWD) in Figures S14 **C** and **D**. The conclusion still holds for each glycan using this metric. However, as cautioned by Bresnett et al. [34], the scoring is quite sensitive to outliers, especially for proper dihedrals with non-Gaussian distributions where unmatched long tails can inflate the score. Therefore, nWD was not used as the primary metric in this work.

### Generic three-monomer angular potentials improve the structural agreements

To assess the structural properties of the generic CG models, we compared the solvent-accessible surface area (SASA) and radius of gyration (*R_g_*) with the AA references. As shown in Table S3, the generic models, denoted as *genNo*, showed moderate agreement with the AA reference in terms of average SASA. The CG models generally produced slightly lower SASA values, reflected by negative percentage deviations. The discrepancies remained below 10%, except for M9, where the deviation was observed -11.0%. Meanwhile, from Table S4, the *R_g_* values of the *genNo* models showed inconsistent agreement with the AA reference, where smaller glycans such as M5 and NGA2F only deviated by less than 10% from the AA models, while largers glycans such as A2 and F1N4 showed worse agreement (*>* 10%). The discrepancy also increased for intermediate-sized glycans, with *R_g_* values reduced by more than 5%. It is clear that the current GMT *genNo* models failed to reproduce the AA *R_g_* distributions, yielding artificially compact conformations.

Further, analysis of branch-distance distributions, defined as the distance between the core *β*-mannose (Man*β*(1 *→* 4)GlcNAc*β*(1 *→* 4)GlcNAc) and the terminal monosaccharide of each branch, the poor agreement primarily resulted from excessive branch flexibility. As shown in Figure S15, the *genNo* models generally sampled shorter and more broadly distributed branch distances relative to the AA references. This reduced branch rigidity largely contributed to the underestimated *R_g_* distributions.

Previous Martini 3 carbohydrate work [12] showed that adding dihedrals spanning four beads across three consecutive monomers may improve the rigidity. That was demonstrated for homogeneous carbohydrate polymers such as dextran and cellulose, where repeating linkage patterns allow the consistent propagation of the dihedrals. However, complex heteropolysaccharides, such as in our N-glycans, are challenging due to their heterogeneous linkages and multi-branched topology. Threemonomer dihedral terms increase the risk of numerical instability because of the overlapping dihedral potentials with the existing two-monomer linkage dihedrals. This issue is particularly pronounced in the branching points, as discussed in the previous section, where multiple dihedrals may act on shared beads. Therefore, three-monomer dihedrals are less straightforward within our building-block parametrization framework.

We instead introduced the three-monomer angle, where the virtual site of the middle monomer acts as a vertex. This construction can be generalized for two different three-monomer configurations, as shown in Figure 4**A**. The three-monomer dihedral configuration proposed by the previous work [12] only covers the top configuration. However, defining an angle for the bottom configuration is more straightforward than the dihedral. The latter configuration occurs in the Man(*α*1 *−* 2)Man units of high-mannose glycans. We posited that these angular terms provide “soft constraints” to increase branch rigidity.

**Figure 4:**
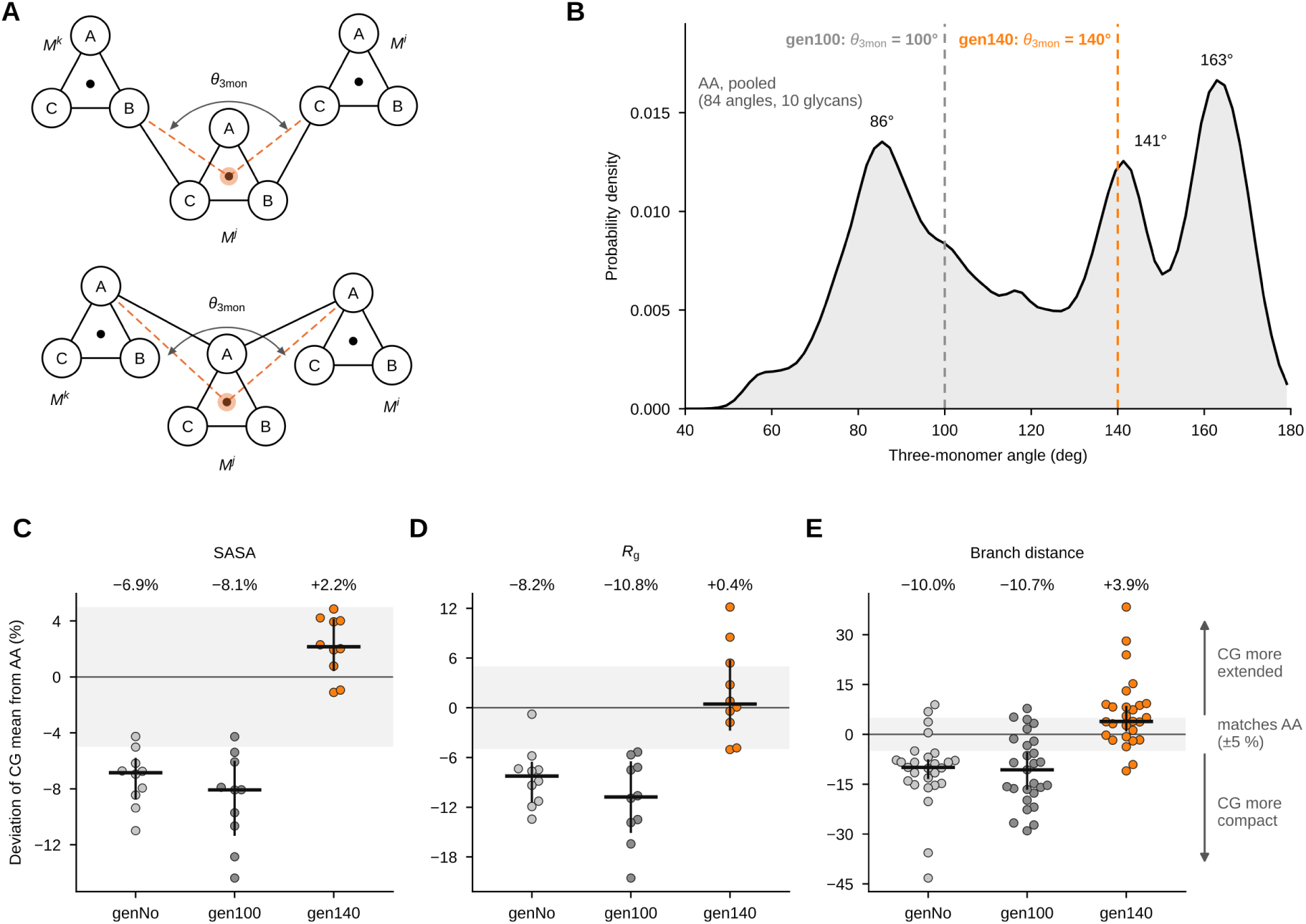
A generic three-monomer angle restores the global shape of the GMT glycans.A. The three-monomer angle *θ*_3mon_, with its vertex at the virtual site of the central monomer *M^j^* (orange) and its arms to the neighbouring monomers *M^k^* and *M^i^* (dashed orange lines), defined for the two bead configurations that occur in the glycan panel (top, B–C links; bottom, A–A links, as in Man*α*(1 2)Man). **B.** Pooled distribution of the three-monomer angles in the mapped AA trajectories (84 angles along the glycan trees of the ten glycans; angles between branches and angles involving fucose excluded), with peaks at 86°, 141° and 163°. Dashed lines mark the tested values: 100° (*gen100*, grey) and 140°(*gen140*, orange). **C–E.** Deviation of the CG mean from the mapped AA mean, 100 (*µ*_CG_ *µ*_AA_)*/µ*_AA_, for SASA (C), *R_g_* (D) and branch distances (E), without a threemonomer angle (*genNo*, light grey) and with *θ*_3mon_ = 100° (*gen100*, dark grey) or 140° (*gen140*, orange). Points: glycans (C, D; *n* = 10) or branch distances (E; *n* = 27). Bars: median (value above) with 95% bootstrap CI. Grey band: 5%. Negative values: CG more compact. M5 is shown with *gen140*; the final GMT model uses *genNo* for M5.

To parametrize the angles, we again employed a modular strategy. We extracted the three-monomer angles along the glycan tree of each glycan from the mapped AA trajectories (84 angles from the ten glycans; angles between two branches and angles involving fucose were excluded) and pooled them. The pooled distribution is broad and has three peaks, at 86°, 141° and 163° (Fig. 4**B**). The 163°peak comes mainly from extended segments of the antennae and the chitobiose core, whereas the 86°and 141° peaks combine linear and branch-point triads (Supplementary Fig. S13), so no single value represents all glycan regions.

Despite these considerations, we leveraged a parsimonious approach, in which a single ReB potential (see Equation 3, type 10 potential in GROMACS 2025.3) with a constant reference value of *θ*_3mon_ is applied across all glycan regions. Therefore, *θ*_3mon_ needed to be chosen to best preserve the structural agreement with the AA references. Using three exploratory GMT configurations: no three-monomer angle (*genNo*) as a control, a 100*^o^* angle (*gen100*), and a 140*^o^* angle (*gen140*), we utilized three structural descriptors, namely SASA, *R_g_*, and branch distances, to assess each configuration. We tested two reference values, 100° and 140°. 140°coincides with the 141° peak, the mode most populated by branch-point triads, where the *genNo* models lack rigidity (Supplementary Fig. S13); larger values, including the 163° peak, caused occasional numerical instabilities in preliminary 1 *µ*s CG simulations for most glycans.

We then compared the simulation results for the three configurations through *R_g_* and SASA, shown in Figure 4**C** and **D**. Overall, *gen140* increased SASA and *R_g_* relative to *genNo* and *gen100* and brought both close to the AA reference (median deviation: SASA *−*6.9, *−*8.1 and +2.2%; *R_g_ −*8.2, *−*10.8 and +0.4% for *genNo*, *gen100* and *gen140*; Fig. 4**C**, **D**). With *gen140*, the SASA of all ten glycans was within 5% of the AA reference, compared with one glycan for *genNo* (Table S3). M5 glycan still showed excellent agreement in both *genNo* and *gen140* strategies. However, the 100*^o^* angle slightly worsened the SASA agreement compared to the *genNo* result, especially for the larger glycan, F1N4, with a deviation of more than 10% from the AA reference. Likewise, *gen100* also worsened the *R_g_* agreement, which generally increased the deviation to 10% from the AA reference compared to the *genNo* configuration for intermediate-and large-sized glycans. In contrast, the *gen140* configuration improved the agreement, reducing the discrepancy by around 50% relative to the *genNo* for most glycans (see Table S4). For larger glycans, such as A2 and F1N4, the CG gen140 model better preserved extended conformations, with deviations from the AA reference below 10%. Additionally, it is noted that M5 was the only glycan for which adopting *gen140* worsened the agreement, where the discrepancy was +12.2%. In contrast, the discrepancy of M5 with *genNo* was only *−*0.8%. We therefore opted to use *genNo* for M5 and *gen140* for all other glycans examined.

Locally, the *R_g_* trend follows directly from the branch distance distributions. More specifically, the *gen140* configuration for the largest glycan in our panel, F1N4, showed moderate agreement with the AA reference for the four branch distance distributions compared to *genNo* setup, as demonstrated in Supporting Figure S18. On the contrary, the *gen100* configuration yielded lower branch distances, leading to poor *R_g_* agreement. Across all 27 branch distances (Fig. 4**E**), *gen140* deviated least from the AA reference (median +3.9%, 12 of 27 within *±*5%), whereas *genNo* and *gen100* gave systematically shorter branches (median *−*10.0% and *−*10.7%; 2 and 6 of 27 within *±*5%). From this, we concluded that a generic three-monomer ReB angle potential with *θ*_3mon_ = 140*^o^* was sufficient to refine the rigidity of all glycans except M5 in the GMT force fields.

To assess the transferability of the *gen140* strategy, we employed G2F2N1, a complex N-glycan outside of the parametrization panel (Figure 5**A**). Its final CG topology, shown in Figure 5**B**, was assembled using the building block approach, following its SNFG representation in Figure 5**A**. All linkage unit parameters were available from the generic linkage library. Next, we performed the 3-*µ*s CG simulation and compared the structural properties with its corresponding 3-*µ*s AA reference simulation. The resulting : SASA, *R_g_*, and branch distance distributions are shown in Figures 5 **C,D**, and **E**, respectively. SASA and *R_g_* of the CG model show reasonable consistency with the AA reference, albeit with generally broader distributions. The discrepancy found in the branch distance distribution for the longest branch (Br. 1) may contribute to the broader *R_g_* distribution, despite a good agreement for other branches.

**Figure 5:**
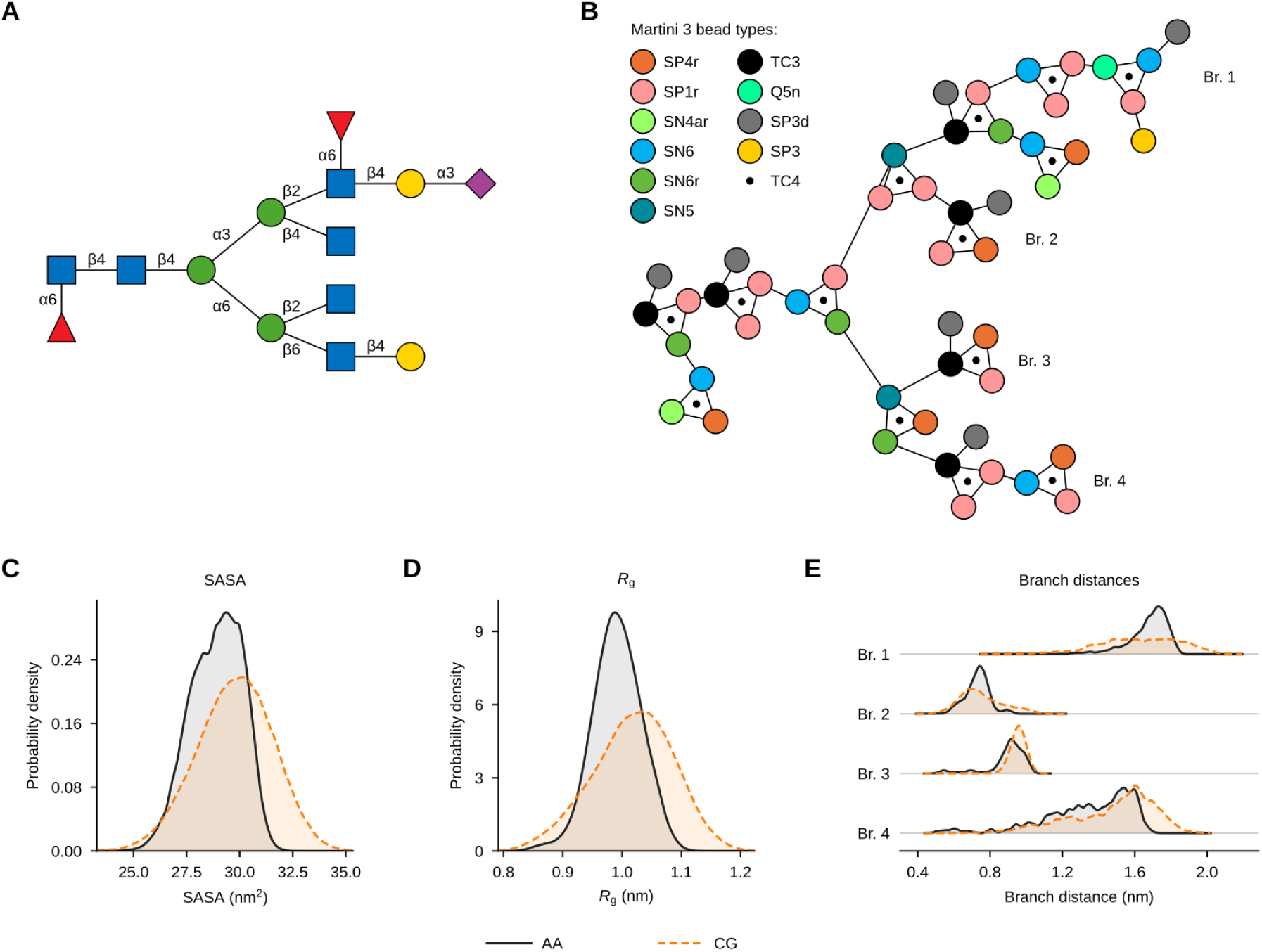
Transferability of the GMT model to G2F2N1, a glycan outside the parametrization panel. **A.** SNFG of G2F2N1 (GlycoSHIELD entry Gal2 Fuc2 Neu1), with a core *α*(1 6) fucose and a second *α*(1 6) fucose on the GlcNAc of the sialylated antenna, as in the reference structure. **B.** GMT model assembled from the linkage library, with beads coloured by type. Br. 1–4: branch distances shown in E, from the A bead of the core *β*-Man to the A bead of the terminal residue of each branch. **C–E.** SASA (C), *R_g_* (D) and branch distances (E) from the mapped AA (black, grey fill) and CG (orange, dashed) simulations. Curves are smoothed for display; in E, each AA/CG pair is scaled by a common factor.

### Glycan-specific parametrization benchmarks the generic transferable model

In this section, we described the non-modular framework, GMR strategy, independently optimizing linkage parameters for each glycan topology without pooling shared linkage units or assuming transferability. This molecule-specific approach, commonly used in the Martini force field development, aimed to benchmark our proposed generic models. Additionally, restricted bending (ReB) angle potentials were manually parametrized for each adjacent monosaccharide triad.

The final GMR glycan linkage parameter set (*CG Manual*) was compiled and compared against the CG *gen140* model in Figures S3–S12. This optimized set produced satisfactory agreement between the CG and AA distributions. Further assessment using the FF and nWD scores further supported this agreement (Figure S19), with the total FF score remaining consistently below 0.3 and the total nWD below 0.5.

Overall, the two parametrization strategies also showed comparable SASA and *R_g_* distributions (Fig. S16**A** and **B**; Table S5). The manually optimized GMR models reproduced AA references with both SASA and *R_g_* deviations below 5%. Relative to the GMR models, the SASA for *gen140* models deviated by less than 5%, whereas *R_g_* deviated below 10% across all glycan sizes. Pooled branchdistance distributions from the ten glycans also showed a moderate positive average increase (+6.2%) compared to the other strategies. These results further support generic parametrization, *gen140* as a practical approximation to the CG construction of moderate-sized N-glycans.

### 2.4 Parametrization of the asparagine–glycan linkage in the N-glycoproteins

To implement the glycan parameters for the N-glycoprotein construction, we defined the CG covalent link between the asparagine side chain and the reducing end of GlcNAc, termed GlcNAc–Asn linkage. The linkage comprises one bond, three angles, and four dihedrals involving the preceding (*−*BB) and following (+BB) backbone beads, the Asn BB and SC1 beads, and beads from the first and second GlcNAc residues (Fig. 6**A**). Two dihedrals (*−*BB, BB, SC1, A1) and (+BB, BB, SC1, A1) were introduced to maintain the upright orientation of the glycan relative to the tripeptide backbone, whereas (BB, SC1, A1, C1) controls GlcNAc torsion around the Asn side chain. The additional dihedral (SC1, A1, B1, A2) controls the flipping of the first GlcNAc aromatic ring. The CG parameters were derived from pseudo-CG (AA) trajectories of ten systems of tripeptide (Gly-Asn-Gly) + N-glycan in the GlycoSHIELD database [35]. Following the modular strategy, linkage parameters were obtained by averaging the pooled distributions from the ten tripeptide systems and fitting each distribution to its corresponding bonded interaction potential. The final parameter set was reported in Table S6.

**Figure 6:**
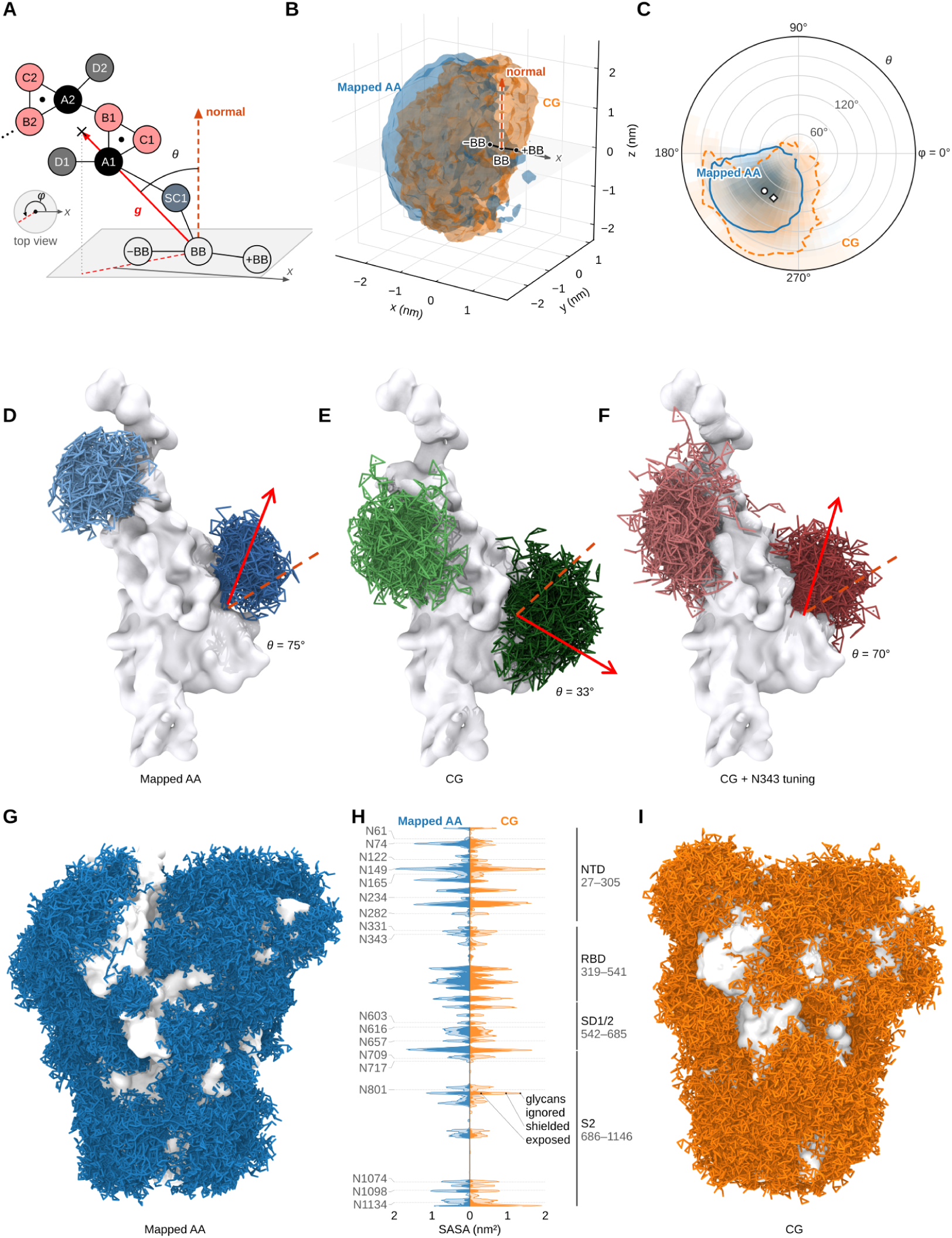
GlcNAc–Asn linkage and glycan orientation in a tripeptide, the RBD and the full SARS-CoV-2 spike. **A.** Beads defining the GlcNAc–Asn linkage ( BB, BB, +BB, Asn SC1, GlcNAc beads A1–D1, A2–D2; dots, virtual ring centres) and the orientation angles: *θ*, between the backbone normal (red-orange, dashed) and the glycan vector *g*; *φ*, azimuth of *g* from *x* (top view; Methods). **B, C.** M9 on Gly–Asn–Gly, mapped AA (blue) vs CG (orange): 80% occupancy surfaces (B) and angular map with *θ* as radius and *φ* as polar angle (C); markers, mean glycan directions (AA 78.0°/227.6°, CG 78.4°/240.6°; 12.8° apart). **D–F.** RBD with M5 at N331 (light) and N343 (dark): mapped AA (D), generic CG (E), CG with N343 tuning (F); *θ* of the mean glycan direction at N343; arrows are schematic. **G–I.** Glycosylated spike trimer, generic CG model (4 replicas, 10 µs): glycan conformers from mapped-AA (G, blue) and CG (I, orange) trajectories, 80 per site; protein surface white. **H.** Per-residue SASA (chain average), mapped AA left, CG right: exposed, shielded by glycans, and glycans ignored; dotted lines, glycosites; bars, domains.

Using the tripeptide-M9 system, we first compared the resulting GlcNAc-Asn linkage distributions with the AA reference, shown in Figure S20. The CG model showed reasonable consistency across the linkage parameters. To further assess the spatial distribution and the orientation of the attached glycan, we computed three-dimensional occupancy maps from both mapped-AA and CG trajectories, where the peptide backbones served as the anchor. The resulting maps, shown in Figure 6**B**, showed a correct cloud alignment and substantial overlap between the AA and CG glycan clouds. Moreover, the orientation preference was quantified by representing the 3D occupancies as a two-dimensional polar plot in Figure 6**C**, where the angular coordinate describes the glycan rotation around the backboneplane normal, and the radial coordinate is the tilt angle relative to this normal. Although the CG model sampled a broader angular range, the mean tilt angles were nearly identical between AA and CG (77.98° and 78.43°, respectively), as well as the 3D angular separation of only 12.79°.

### 2.5 GlycoMarinate applications on glycoprotein systems

#### Site-specific evaluation from CG simulations of receptor binding domain of SARS-CoV-2 spike protein

By integrating both the glycan and GlcNAc-Asn linkage parameters into the GlycoMarinate workflow (see Figure 1), we examined the transferability of the proposed parametrizations in a full protein context. For the first case, we utilized a receptor-binding domain (RBD) taken from the full SARSCoV-2 spike protein in upward conformation, described in Yang et al. (2021) [36]. The domain contains 229 residues (aa 319-547) with two known N-glycosylation sites: Asn331 and Asn343 (see Figure S21 **A**). The sites represent two contrasting local environments, where Asn331 is located in a highly flexible long loop while Asn343 is in the short loop adjacent to the antiparallel *β*-sheet. We prepared a simple system bearing two M5 glycans and performed 2*µ*s CG simulations alongside corresponding AA simulations using the CHARMM36 force field.

Focusing on the two tripeptide segments (Pro330-Asn331-Ile332 and Phe342-Asn343-Ala344), the pseudo-CG trajectories mapped from the AA simulation were aligned with the CG trajectories. The resulting distributions at each glycosylation site were compared with the AA RBD simulations, shown in Figure S22 **A** and **B**. Overall, the CG angle distributions reasonably reproduced the AA references, with peak deviations generally below 30°. A site-dependent difference occurred for the (SC1, A1) bond, whose AA peaks differed by 2.5 Å between Asn331 and Asn343. Although the generic CG parameters did not capture this shift, the broader CG distribution still encompassed both AA peaks. At Asn331, all four dihedral distributions showed a reasonable alignment with the AA reference. On the contrary, at Asn343, the (BB, SC1, A1, C1) dihedral failed to reproduce the distinct AA peak near *−*27°.

To evaluate how the altered dihedrals affect the orientation of the whole glycan cloud, we projected the glycan occupancy onto the *xy* plane of the local backbone frame (Supplementary Fig. S23**A**). At Asn331, the CG cloud pointed in the same direction as in the AA simulation, with the peaks of the in-plane orientation within 20° (Supplementary Fig. S23**C**). At Asn343, by contrast, the CG glycan stood more upright: its mean direction made an angle *θ* = 33° with the backbone normal, compared with 75° in the AA reference (azimuth *φ* = 265° vs 220°; Fig. 6**D**, **E**; Methods). Of the four linkage dihedrals, (BB, SC1, A1, C1) and (SC1, A1, B1, A2) controlled the overall cloud orientation.

We therefore refitted these two dihedrals at Asn343 to the mapped AA distributions of the glycosite tripeptide (Phe342-Asn343-Ala344) in the RBD simulation, which moved the minimum of the (BB, SC1, A1, C1) dihedral from 185° to *−*20°, close to the AA peak at *−*27°, and that of the (SC1, A1, B1, A2) dihedral from 42° to 90°, with a stiffer (BB, SC1, A1, C1) term (*k_ϕ_* 12 to 20 kJ mol*^−^*^1^; Supplementary Table S6). This improved the CG dihedral distributions (Fig. S22**C**). More importantly, the tuning restored the orientation of the Asn343 glycan, giving *θ* = 70° and *φ* = 220° compared with 75° and 220° in the AA reference (Fig. 6**F**; Supplementary Fig. S23**B**, **C**).

Additionally, we also reached the same conclusion when the M5 glycans were replaced with the NGA2F glycans (Fig. 2**C**), in which a fucose moiety is (*α*1 *−* 6)-linked to the first GlcNAc. In brief, we still discovered the site-specific discrepancy for the linkage parameters in the CG simulations (Figure S24). However, the similar tuning from the previous M5 case on dihedrals (BB,SC1,A1,C1) and (SC1,A1,B1,A2) recovered the overall glycan cloud orientation (Figure S25). Hence, the approach is applicable to both high-mannose and complex glycans.

The site-specific behavior may arise from aromatic–carbohydrate interactions between Phe342 and the first GlcNAc ring, potentially restricting ring flipping and the distributions of the two dihedrals. Such effects may be weaker at Asn331, where the nearby aromatic residue Phe329 is located further from the glycosylation site. Following that observation, we therefore propose “soft” tuning for the two glycosidic dihedrals in an aromatic sequon, namely *R*_1_-Asn-*R*_2_-Ser/Thr, where Asn is any glycosite with at least one of (*R*_1_, *R*_2_) being one of the aromatic amino acids (Phenylalanine, Tyrosine, Tryptophan). The conditional dihedral parameters are reported in Supplementary Table S6.

### Glycan shielding of fully glycosylated trimeric spike protein from CG simulations

We also turned to the full SARS-CoV-2 trimeric spike glycoprotein, for which extensive AA simulation data exists [37]. In particular, we only used amino acid residues 27-1146 from each subunit of the ectodomain of the spike protein, retrieved from the 6VSB PDB structure [38]. The atomistic structure was mapped to a CG representation and used as input to the GlycoMarinate pipeline to attach the 18 CG glycans per subunit with the closest composition to that reported in published mass-spectrometry datasets [39] (see Fig. S26). We obtained a 10 *µ*s CG simulation at Δ*t* = 20 fs timestep.

Because surface accessibility of viral spike proteins determines epitope exposure and is therefore central to both natural and vaccine-induced immunity, we next assessed how faithfully the simulations reproduce glycan-mediated shielding of the protein surface. The visualized comparison with the AA-mapped glycan shielding is demonstrated in Figure 6**G–I**. The glycan-induced change in SASA, ΔSASA, showed good agreement with the AA-mapped reference simulation when evaluated using a 0.191-nm probe radius. The resulting site-resolved shielding profiles correlated well with those obtained from AA simulations (Pearson’s *r* = 0.76; Fig. 6**H**), with comparable mean SASA values (0.42 vs. 0.39 nm^2^ per residue). Both models captured similar relative exposure of the key domains: NTD, RBD, SD1/2 and S2. Using the glycan shielding score definition: ΔSASA = (SASA_nogly_ *−* SASA_gly_)*/*SASA_nogly_), the RBD is the least occluded region (*<* 14%), consistent with its epitope exposure property. A closer fraction agreement was shown for SD1/2 region with 17% and 19% for the CG and AA models, respectively. However, some regions in the CG model overestimated the shielded fraction, such as S2 (24% vs 14%) and NTD (23% vs 17%). Shielding overestimation is expected in CG-level glycans due to the large effective bead radii. Overall, the results indicate that GlycoMarinate-enabled CG simulations can reproduce biologically relevant patterns of glycan shielding.

## 3 Methods

### 3.1 Simulation engine

Unless indicated otherwise, simulations were performed using GROMACS 2025.3 [40] or newer, including related analysis modules.

### 3.2 Reference all-atom simulations

#### Simulations of N-glycans

Atomistic reference trajectories of the ten glycopeptides described in the panel (Fig. 2**D**), including the G2F2N1 system (Figure 5**A**), were retrieved from the GlycoSHIELD conformer library [35], which contains 3 *µs* trajectories of 68 N-and O-glycans. The simulation preparations and production runs are detailed by Tsai et al. (2024) [26]. In brief, glycan structures were generated using the averaged fragments approach implemented via Glycan Modeler [24], while CHARMM-GUI [41, 42] was used for generation of solvated glycopeptides, where the glycan was attached to a central asparagine residue of a Gly–Asn–Gly tripeptide. All systems used CHARMM36m force field [30, 31, 43]. The resulting trajectories contain at least 3 *µs*. In this dataset generation, GROMACS 2018.0 was used in all production runs.

#### All-atom simulations of glycoproteins

First, we retrieved the atomistic structure of the receptor-binding domain (RBD) of the SARS-CoV-2 spike (S) protein from the cryo-EM structure of a full S protein in “RBD-up” conformation (PDB ID: 7EAZ) [36]. The domain consists of 228 residues (i.e., aa 319-547 from the S protein). We prepared the simulation input files using CHARMM-GUI, where the glycans at Asn331 and Asn343 were remodelled with Glycan Modeller [24]. The atomistic CHARMM36m force fields were selected for all systems. Two glycoproteins were prepared in which the two sites were attached with either M5 or NGA2F glycans. Each solute was dissolved in a dodecahedral system filled with TIP3P water molecules [44]. The systems were then neutralized with a NaCl concentration of 150 mM. After standard energy minimization and equilibration, two coupling algorithms: the velocity rescaling (coupling temperature with *τ* = 1 ps) [45] and Parrinello-Rahman (coupling pressure with *τ* = 1 ps and *β* = 4.5 *×* 10*^−^*^5^ bar*^−^*^1^) [46] were utilized for all production runs to retain a constant temperature of 310 K and pressure of 1 atm. All simulations were run for 2*µ*s with Δ*t* = 2 fs.

##### Spike glycoprotein: atomistic reference

As the atomistic reference for glycan shielding, we reanalysed the published 2.5 *µ*s CHARMM36m trajectories of four fully glycosylated SARS-CoV-2 spike trimers in a membrane [37] (PDB 6VSB; glycoforms from site-specific glycoproteomics [39]; simulation lengths in Table 3). Each trimer was analysed separately, and only the ectodomain (residues 27–1146) was used, which leaves 18 of the 22 N-glycosylation sites per protomer (N17, N1158, N1173 and N1194 lie outside this range). The glycans at these sites are shown in Fig. S26.

**Table 3:** Summary of simulation sampling. Replica counts and durations apply separately to each system at each resolution. Each spike protein system was simulated using four independent replicas of 2.5 *µ*s at each resolution. All other systems used a single run.

| System or purpose | Resolution | Replicas | Simulation time (*µ*s) |  |
|---|---|---|---|---|
|  |  |  | Per replica | Total |
| Glycosidic dihedral screening | CG | 1 | 1 | 1 |
| 10 glycopeptides | AA and CG | 1 | 3 | 3 |
| A2 and M9 aggregation | AA and CG | 1 | 1 | 1 |
| Single G2F2N1 simulation | AA and CG | 1 | 3 | 3 |
| RBD protein systems | AA and CG | 1 | 2 | 2 |
| Spike protein systems | AA and CG | 4 | 2.5 | 10 |

### 3.3 Bonded interaction parametrizations

At the monosaccharide level, we retained the original internal bonded parameters (intramonomer bonds, angles, and dihedrals) inherited from the Martini 3 carbohydrate model. In the Martini 3 carbohydrate parametrization, the constraint lengths were obtained from AA reference simulations. The constraint lengths of all sugar rings were uniformly increased by 15% to compensate for the molecular volume underestimation obtained from direct CoG mapping [12].

Therefore, we only focused our CG parametrization on bonded interactions introduced by glycosidic linkages and the GlcNAc–Asn connection. The bonds between two linkage beads are parametrized by reference bond length and force constant, denoted by *r*_ref_ and *k_r_*, respectively, within a harmonic potential function:

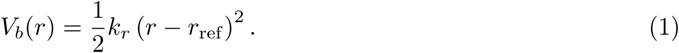

The harmonic potential is also used to parametrize two linkage angle potentials:

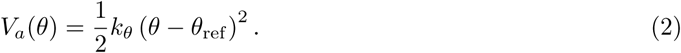

The reference values and force constants are denoted by *θ*_ref_ and *k_θ_*. On the other hand, the restricted bending (ReB) potential is applied to avoid numerical instability when the formed angle becomes collinear [47]:

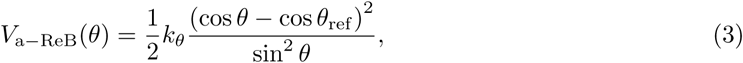

In this work, this potential is generally applied if the angle distributions have a maximum located above 135*^o^*. Next, for all proper dihedrals, we applied a periodic function:

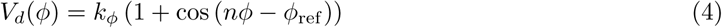

where *k_ϕ_*, *ϕ*_ref_, and *n* refer to the force constant, reference angle, and multiplicity, respectively. For simplicity in generic parametrization, we assumed that a single-minimum dihedral, *n* = 1, is sufficient for parametrizing all linkage units. In addition, the improper dihedrals are used only for retaining intramonomer dihedrals, especially original chirality in GlcNAc and Neu5Ac:

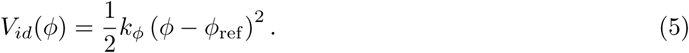

To extract the corresponding linkage parameters from the all-atom reference simulations, CG mapping was first performed on each trajectory frame to yield pseudo-CG trajectories. The mapping strategy utilized the atom-to-bead “dictionary” proposed in S1. Initial equilibrium values and force constants for the bonded potentials were obtained from the corresponding AA distributions, extracted using the Boltzmann inversion (BI) method. For each bonded coordinate (x), the effective potential was calculated as (*U* (*x*) = *−k*_B_*T* ln *P*_AA_(*x*)), where (*P*_AA_(*x*)) is the probability distribution obtained from the mapped AA ensemble. The set of parameters was then determined by fitting the resulting effective potential to the functional form used for the corresponding Martini bonded interactions. The trajectory analyses and BI fitting were implemented using an in-house script by utilizing MDAnalysis, Numpy, and Scipy packages in Python 3.12.

In the generic parametrizations, the **GMT** strategy used the final fitting parameters from the cumulative/pooled AA distributions across multiple glycans directly, without further tuning. In contrast, for the second parametrization strategy, **GMR**, the fitting parameters were used to construct the initial CG topology for the first CG simulations. The resulting bonded distributions were investigated directly with the corresponding AA reference distributions to identify parameters requiring further refinement. The parameter tuning was iterated until the CG distributions closely matched the AA distributions.

### 3.4 Coarse-grained simulations

#### Simulations of N-glycans

CG glycopeptide models were generated from atomistic reference structures obtained from the GlycoSHIELD library. Each structure was mapped to the CG resolution using *martinize2* and also the atom-to-bead mapping scheme described above. The systems were put into the center of the cubic boxes with a minimum distance of 3.5–5.0 nm to the box edge, depending on the glycan sizes. The systems were solvated with Martini water and neutralized with sodium or chloride ions, with a total salt concentration of 150 mM. The energy minimization was then conducted and continued with equilibration for 125 ps in the NVT ensemble and 125 ps in the NPT ensemble at 300 K using the velocity-rescaling thermostat with a coupling constant of 1.0 ps. During NPT equilibration, pressure was retained isotropically at 1 bar using the Berendsen barostat with (*τ_p_* = 5.0) ps and a compressibility of (3 *×* 10*^−^*^4^)bar(*^−^*^1^). Moreover, nonbonded interactions were set with the Verlet cutoff scheme using a cutoff of 1.1 nm for both Coulomb and Lennard-Jones interactions, with potential-shift modifiers and a relative dielectric constant of 15. Finally, the production simulations were run for 3 *µs* for each system (see Table 3) with a 20 fs integration time step. The resulting trajectories were aligned to the corresponding atomistic reference using the tripeptide backbone beads.

#### Simulations of glycoproteins

CG structures and topologies of the spike trimer and of the RBD were generated from the atomistic starting structures with martinize2 (Vermouth 0.13) [48], using Martini 3.0.0 [8] and the GōMartini 3 model [21] instead of an elastic network. Secondary structure was assigned by martinize2 (DSSP), and Gō contacts were derived from the same structures (for the spike, the 6VSB-based atomistic model) with contact cut-offs of 0.3–1.1 nm, a minimum sequence separation of three residues, and a well depth of *ε* = 9.414 kJ mol*^−^*^1^. For the spike, the three protomers were merged into one molecule, so the 6,134 contacts include 467 between protomers. Disulfide bonds were detected automatically, and termini were kept neutral.

The glycoprotein models were then built using the GlycoMarinate pipeline based on the provided glycosylation list. For the RBD, following the corresponding AA simulations, we prepared two systems: RBD with two M5 glycans and RBD with two NGA2F glycans. For the trimeric spike protein, CG glycans were attached at the 18 glycosylation sites per protomer (Figure S26). They were chosen from our parametrized panel (see Figure 2**D**) to match the atomistic glycoforms. The panel had no exact match at five sites, where we used the closest glycan: NA2 instead of the core-fucosylated FA2G2 at N149, N343, N616 and N1134, and Man9 instead of Man8 at N234. At the four sialylated sites (N74 and N331 with G3F1N1, N165 and N1098 with A1F), the CG glycans carry Neu5Ac*α*(2*→*6)Gal, whereas the atomistic model has *α*(2*→*3). Because the trimer is densely glycosylated, we raised the GlycoSHIELD clash cutoff from the default 0.7 nm to 0.8 nm [26].

For each system, the protein was placed in a cubic simulation box with solute–box buffer distances of 4.5 nm and 5.1 nm for the RBD-only and the full spike systems, respectively, followed by an initial energy minimization in vacuum. We then solvated each minimized structure with equilibrated Martini water and neutralized it with 100 mM NaCl, followed by a second energy minimization. To allow the solvent, ions, protein, and glycans to equilibrate due to the initial restrained glycan-protein contacts, the system was equilibrated gradually through multi-step NVT and NPT simulations using gradually increased integration time steps. V-rescaling thermostat (*τ* = 1 ps) and a Parrinello-Rahman barostat (*τ* = 12 ps and *β* = 3 *×* 10*^−^*^4^ bar*^−^*^1^) were applied to keep the temperature constant at 300 K and the pressure at 1 atm during the total 10*µs* production simulations (see Table 3). Integration timesteps of 20 fs were used for all production runs.

### 3.5 All-atom and coarse-grained agreement

The agreement between CG and AA reference distributions was quantified using the normalized first Wasserstein distance [49] (nWD). For reference AA (*P*) and fitted CG (*Q*) distributions, the Wasserstein distance was calculated as 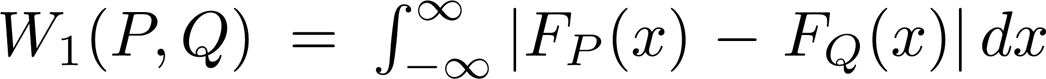, where *F_p_* and *F_Q_* are the corresponding empirical cumulative distribution functions. To express deviations relative to the intrinsic width of the atomistic distribution, *W*_1_ was normalized by its interquartile range (IQR), such that nWD = *W*_1_(*P, Q*)*/*IQR_AA_, with IQR_AA_ = *Q*_3_ *− Q*_1_.

Agreement was further quantified using the Fast-Forward score as defined in Brasnett et al. [34]. For normalized AA and CG probability distributions *p* and *q*, respectively, the score was calculated as

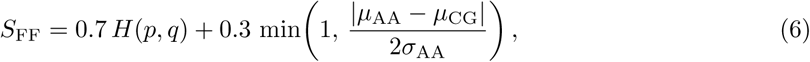

where

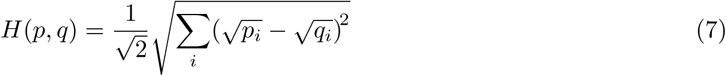

is the Hellinger [50] distance, *µ*_AA_ and *µ*_CG_ are the respective distribution means, and *σ*_AA_ is the standard deviation of the AA distribution. The mean-displacement contribution is capped at unity. Following the Fast-Forward score convention, values below 0.3 indicate good agreement, values between 0.3 and 0.5 indicate acceptable agreement, and values above 0.5 indicate poor agreement [34]. Both nWD and *S*_FF_ scores were calculated using in-house Python scripts for bonded linkage distributions, including bonds, angles, and dihedral angles. Dihedral angle distributions were treated using circular statistics.

### 3.6 Molecular shape and volume agreements

Solvent-accessible surface area (SASA) was calculated using the double cubic lattice method [51] implemented in GROMACS 2025.3 via gmx sasa command. Here, the suggested Martini VdW radii for different bead sizes were used: 0.264 nm for regular beads, 0.230 nm for small beads, and 0.191 nm for tiny beads. The probe size is 0.191 nm. Meanwhile, the radii of gyration (*R_g_*) of AA and CG glycans were also computed in GROMACS using the gmx gyrate command. Branch distances were calculated frame by frame as the distance between the A bead of the core *β*-mannose and the A bead of the terminal monosaccharide of each branch, using the CG trajectories and the mapped AA (pseudo-CG) trajectories, giving 27 branch distances for the ten glycans (2–4 per glycan) and four for G2F2N1. Branch distances were calculated frame by frame as the distance between the A bead of the core *β*-mannose and the A bead of the terminal monosaccharide of each branch, using the CG trajectories and the mapped AA (pseudo-CG) trajectories, giving 27 branch distances for the ten glycans (2–4 per glycan) and four for G2F2N1. For each CG setup, agreement with the AA reference was expressed as the signed deviation of the CG mean from the AA mean, 100 (*µ*_CG_ *− µ*_AA_)*/µ*_AA_, per glycan (SASA, *R_g_*) or per branch distance, and summarised as the median with a 95% percentile bootstrap confidence interval (10,000 resamples of the glycans or branch distances).

### 3.7 Occupancy map

To make a visual comparison of the spread and spatial sampling of CG glycans against the reference AA simulations, we computed the occupancy density maps of glycans linked to tripeptides. First, the backbone beads of the CG trajectories were aligned to the backbone atoms of their AA counterparts. From this, we created a bounding box around the systems using the maximum and minimum glycan positions, with a 10 Angstrom padding. The occupancy of atoms or beads over the trajectory was then computed within the box using the DensityAnalysis routine in MDAnalysis with three-dimensional grids of 1 Å.

### 3.8 Glycan shielding score

Glycan-mediated shielding was quantified as the relative change in per-residue solvent-accessible surface area: ΔSASA = (SASA_nogly_ *−* SASA_gly_)*/*SASA_nogly_), following the shielding score definition in the GlycoSHIELD procedure [26]. For glycan ensembles, SASA_gly_ was normalized over all conformations, such that ΔSASA = 0 indicates no glycan shielding and ΔSASA = 1 complete shielding of the residue. Shielding of the SARS-CoV-2 spike protein was calculated using a probe radius of 0.191 nm. Moreover, as a direct comparison with the atomistic system, the time-dependent shielding score for the AA simulation was calculated on the pseudo-CG trajectories. The atom-to-bead mapping for glycan parts was taken from the individual glycan mapping, whereas the default amino acid mappings between atomistic CHARMM and coarse-grained Martini 3 force fields were used for the protein part [8, 48]. Before mapping, glycans in the atomistic trajectories were made whole across the periodic boundaries by walking the bond network outward from the protein. For the visual comparison of glycan clouds (see Figure 6**G**-**I**), each trimer was centered and aligned on its S2 core.

#### 3.8.1 Aggregation study

We also investigated the aggregation trends of high-mannose and complex N-glycans, represented here by M9 and A2 glycans, respectively. To achieve the desired concentration of 50 g/L, as suggested in [52, 53], we simulated 40 M9 or 30 A2 molecules in a cube water box with adjusted lengths. We first prepared the initial structure of each system at AA resolution and mapped the whole system to CG resolution. For each system, we performed 1 *µ*s CHARMM36 AA with TIP3 water and Martini 3 CG simulations. We then quantified aggregation using network analysis, following the computation proposed by Brandner et al. (2024) for disaccharide systems [52], where, for each simulation frame, glycan molecules were treated as nodes and connected when the separation distance was *<*4.7 Å. We then identified the resulting connected graphs as aggregates.

## 4 Discussion

We described a modular strategy to systematically parameterise ten biologically relevant N-glycans in the Martini 3 force field, along with their linkage to asparagine side chains. The generic CG linkage library underlying the GMT model, composed of 12 glycosidic linkage units, reproduced both local bonded distributions and global structural metrics (SASA, radius of gyration and branch lengths) with accuracy comparable to the glycan-specific GMR models. The library also transferred successfully to G2F2N1, a glycan outside the parameterisation panel. Finally, applied to the fully glycosylated SARS-CoV-2 spike trimer, the framework produced a 10 *µ*s simulation whose site-resolved glycan shielding profile correlated well with atomistic reference data (*r* = 0.76), including the characteristic low shielding of the RBD.

The two levels of description serve different purposes. The GMT model requires no glycan-specific parameterisation: any non-bisecting N-glycan whose linkages are covered by the 12-unit library can be assembled directly from its monomer connectivity, making it the natural choice for glycans without an atomistic reference, for the heterogeneous compositions reported by glycoproteomics, and for large systems in which many distinct glycoforms must be modelled. Its accuracy is adequate for this purpose: all ten GMT models scored within the acceptable FF range against the atomistic reference, and seven within the good range (M5, M9 and A2 at 0.34, 0.33 and 0.31), whereas every GMR model scored within the good range. SASA and *R_g_* of the GMT models deviated from GMR by less than 10% for all glycans, and pooled branch distances deviated from the AA reference by similar amounts in both models (mean +6.2% GMT, +5.6% GMR). The GMR models, in contrast, reproduce the atomistic ensembles more faithfully (SASA within 4%, *R_g_* within 5%) at the cost of per-glycan optimisation against mapped atomistic trajectories. They are therefore preferable when the conformational ensemble of a specific glycan is itself the quantity of interest, for example when quantifying site-specific shielding or glycan– protein contacts, whereas the GMT model is sufficient when glycans act primarily as a steric and electrostatic shield at the protein surface. The two levels are not mutually exclusive: GMR models for additional glycans can be derived by the same procedure and registered in the workflow alongside the generic library, and new linkage units can be added to the GMT library as further atomistic references become available.

GlycoMarinate builds directly on the Martini 3 carbohydrate framework of Grünewald et al. [12], from which we retained the mapping rules, the three-bead ring representation with a central TC4 virtual site, and all intramonomer bonded terms. Our departures concern the glycosidic linkage. First, bead types at the linkage are assigned according to the atoms actually lost on bond formation, including the doubly substituted mannoses at branch points, rather than reused from the isolated monosaccharides. Second, each linkage is described by a single bond, two angles and one dihedral, instead of the four angles and two dihedrals used for linear polysaccharides [12, 17–19]; in branched glycans, where several linkages share beads, the additional dihedrals impose overlapping torsional restraints that we found to destabilise simulations at the branch points. Third, the loss of rigidity that accompanies this reduction is recovered with a three-monomer restricted-bending angle centred on the ring virtual site rather than the three-monomer dihedral proposed for homopolymers such as dextran and cellulose, which is both more robust at branch points and definable for the Man*α*(1 *→* 2)Man geometry that the dihedral does not cover. These choices make the linkage terms transferable across glycan topologies, which is what allows a library of 12 units to assemble the full panel. Finally, previous Martini carbohydrate models stopped at the free glycan; GlycoMarinate adds the parameterised GlcNAc–Asn junction and an automated route from a Martini 3 protein topology and a glycosylation list to a complete, clash-free glycoprotein, which has so far required manual, system-specific construction in Martini 3.

Several limitations remain. The first concerns the excessive conformational flexibility arising from monomer-level linkages in the generic parameterisation, which manifests as broader distributions of radius of gyration and branch-arm distances than in the AA reference. The generic three-monomer ReB angle potential with *θ*_3mon_ = 140° partially circumvents this discrepancy, improving the molecular shape of intermediate-size glycans while yielding only modest improvement for the larger ones (F1N4 and G2F2N1). A test on poly-LacNAc, compared against an AA reference using the CHARMM36 force field, made the origin of the residual discrepancy clearer: its AA *R_g_* distribution was notably narrow, indicating a rigidity that a single three-monomer ReB angle cannot fully capture. Replacing the ReB form with a linear angle potential [54], applied specifically to the longest branch to maintain collinear conformations, better reproduced the narrow distribution. Its general application, however, requires further investigation across longer heteropolysaccharides, with careful evaluation of polymer properties, to ensure that no spurious rigidity is introduced; region-specific elastic-network or Gō-like potentials may also be worth investigating for this purpose. Bisecting GlcNAc and O-glycans fall outside the validated scope of the present library.

The second concerns the protein linkage. The generic tripeptide-derived GlcNAc–Asn junction is a good default but can require site-specific evaluation of the local sequon environment. In the RBD, the generic linkage did not capture the carbohydrate–aromatic interaction between the first GlcNAc and the benzyl side chain of Phe342, which we suspect involves face-to-face or edge-to-face *π* stacking that restricts the torsional states of the glycosidic linkage. The main discrepancies occurred for the Man*α*(1 *→* 6)Man linkage, where three of the four terms differ from the AA reference. The AA bond and the *A*_3_–*C*_3_–*A*_4_ angle are multimodal, whereas the CG model samples a single, slightly longer bond and a single angle between the AA modes. The CG dihedral is also much narrower than the broad, multimodal AA distribution. Only the second angle, *C*_3_–*A*_4_–*B*_4_, is well reproduced. The AA distributions of this linkage vary between glycans, depending on whether the arm carries a *β*(1 *→* 6)-linked GlcNAc or a sialylated antenna or sits next to a fucosylated core. A single generic term therefore represents only the majority state. This site belongs to a class of aromatic-enhanced sequons, AR-(Xxx)*_n_*-Asn-Yyy-Thr, where AR is any aromatic residue, *n* = 1 or 2, and Xxx and Yyy are any residues except proline at Yyy; such sequons adopt distinct reverse *β*-turns that stabilise carbohydrate–aromatic contacts between the aromatic residue, GlcNAc and the threonine [55, 56]. In the spike protein, the only such motif is Asn343 (Phe342-Asn343-Ala344-Thr345), consistent with its narrow dihedral distributions. We therefore provide a conditional dihedral set for aromatic sequons (Supplementary Table S6), and suggest that a general solution will key the linkage parameters to local sequence and secondary structure rather than to a single tripeptide reference.

The third limitation lies in the non-bonded interactions and cannot be addressed by glycanor site-specific bonded terms. Preliminary simulations of 30 glycans in solution indicate that the spurious selfaggregation that affected Martini 2 carbohydrates [16] is substantially reduced but glycan-dependent: M9 remained in dynamic equilibrium between monomers and transient clusters, as in the AA reference, whereas the sialylated complex glycan A2 aggregated irreversibly (Supplementary Fig. S27). A related imbalance is visible at the glycan–protein interface, where the largest deviations from the AA glycan occupancy occur at sites in which glycans sit in deep crevices of the protein surface, consistent with the earlier observation that the TC4 virtual site only weakly improved carbohydrate–aromatic stacking in Martini 3 [12]. Rebalancing carbohydrate non-bonded interactions is ongoing work and is independent of the bonded framework presented here, which will transfer unchanged to updated bead assignments; the osmotic-coefficient matching used to refine protein–sugar interactions in Martini 2.2 [57] offers a direct experimental route for that rebalancing in Martini 3.

The modular design defines a clear route for extension. Because linkage units are parameterised independently of the glycans in which they occur, the library can be expanded systematically from simulations of individual disaccharides, following the approach recently used to build Martini 3 lipopolysaccharides [20], rather than from a curated glycan panel, and refined models for additional glycans can be registered alongside it. The same strategy is expected to extend readily to O-glycans and glycolipids, which share the glycosidic linkage units and differ only in the anchoring term to the protein or lipid. On the workflow side, the monomer-level graph that is currently defined by hand can be generated directly from IUPAC condensed notation or GlyTouCan identifiers, and because coordinate generation accepts any atomistic glycoprotein model as input, structures from CHARMM-GUI, GlycoSHAPE, experimental glycoprotein structures, or glycan-aware structure predictors can be coarse-grained directly, with GlycoSHIELD serving only as a convenient default for clash-free conformer placement. The topology and coordinate steps are built on *martinize2* and *polyply*, so they inherit improvements to those tools and can be incorporated into their ecosystems. Together, these developments make site-specifically glycosylated proteins a routine input for Martini 3 simulations rather than a systemspecific effort, opening glycan-mediated processes such as receptor clustering, immune recognition and viral shielding to study at the scale of membranes, viral particles, and crowded cellular environments.

## Data and code availability

GlycoMarinate is open source and available at https://gitlab.mpcdf.mpg.de/alhidaya/glycomarinate. The repository contains the atom-to-bead mapping dictionary, the generic linkage library (GMT), the glycan-specific parameters (GMR) for the ten N-glycans, the GlcNAc–Asn linkage parameters including the conditional set for aromatic sequons, and a tutorial for building a glycosylated protein. The atomistic glycan reference trajectories are part of the GlycoSHIELD conformer library [35]. The atomistic spike trajectories were taken from ref. [37]. CG trajectories of the RBD and spike systems are available from the corresponding author on request.

## 5 Acknowledgements

AAH, AD, MS: Project financed under Dioscuri, a program initiated by the Max Planck Society, jointly managed with the National Science Centre in Poland, and mutually funded by the Polish Ministry of Education and Science and German Federal Ministry of Education and Research (UMO2021/03/H/NZ1/00003). KM gratefully acknowledges financial support for this research from departmental grant funds (N19/DBS/000023). BF gratefully acknowledges the Max Planck Society for funding. MS work is funded by the National Science Centre. SONATA BIS 14 project, ‘From Static to Dynamic: Integrating Glycan Dynamics into Structural Biology’ (UMO-2024/54/E/NZ1/00439). The authors gratefully acknowledge the support of Gaia AI Factory, the Polish Artificial Intelligence Factory coordinated by the Academic Computer Centre Cyfronet AGH. Gaia AI Factory provides an innovation ecosystem and expert support that facilitate the development and adoption of advanced artificial intelligence technologies. The project is co-funded by the Republic of Poland and the European Union through the EuroHPC Joint Undertaking (EuroHPC JU) under Grant Agreement No. 101314359.

