## Supplementary Information for "Automating Glycosylation of Protein Models in the Martini 3 Force Field with GlycoMarinate"

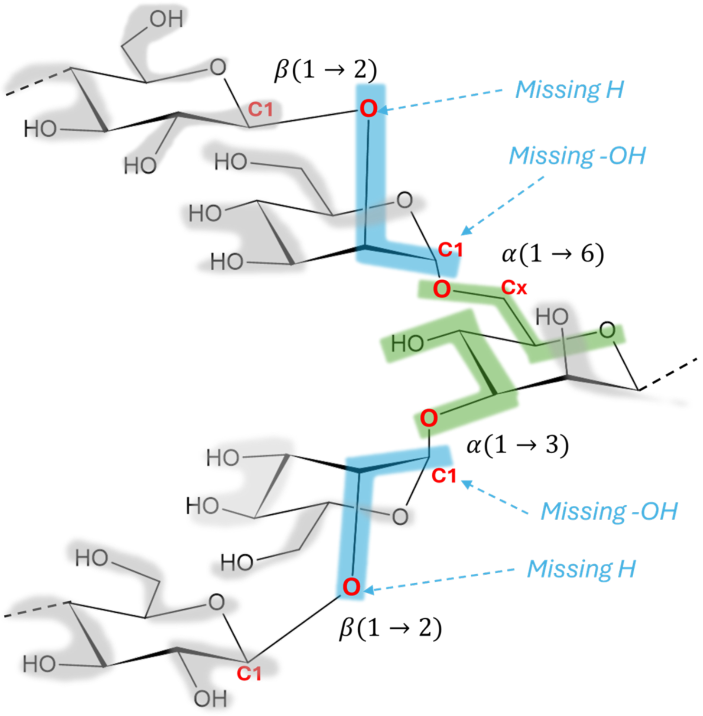


Figure S1: Linkage mapping at the branching point. Two (C$J$–O–C$I$) motifs are highlighted in red. The fragment containing the anomeric carbon (C1) is shaded blue, and the fragment containing the glycosidic oxygen and the acceptor carbon (C$I$) is shaded green. Instead of SN6, the anomeric fragments are assigned SN5 beads, reflecting the reduced polarity caused by the loss of the O2 hydroxyl hydrogen upon GlcNAc$\beta(1 \rightarrow 2)$Man bond formation.

Table S1: Atom-to-bead mapping dictionary used for the Martini 3 carbohydrate representation. Bead names follow the convention $Mx$, where $M$ denotes the bead letter (A–F) and $x$ is the monomer index. For example, the first monomer is represented by beads A1, B1, C1, whereas additional functional groups are denoted by D1, E1, or F1. Linkage-dependent mappings describe beads whose atom composition changes upon glycosidic bond formation.

| **Bead** | **Martini type** | **Atom mapping** | **Application** |
|---|---|---|---|
| A | SP4r | C1, H1, O1, HO1, C2, H2, O2, HO2 | Standard pyranose bead |
| A | SN6 | C1, H1, C2, H2, O2, HO2 | Anomeric linkage bead |
| A | SN5 | C1, H1, C2, H2, O2 | Anomeric bead in $\alpha(1 \rightarrow 3)$ branch |
| A | TC3 | C1, H1, C2, H2 | GlcNAc linkage bead |
| A | Q5n | C1, O11, O12, C2, C3, H31, H32 | Neu5Ac linkage bead at neutral pH |
| B | SP4r | C3, H3, O3, HO3, C4, H4, O4, HO4 | Standard pyranose bead |
| B | SP1r | C3, H3, O3, HO3, C4, H4, O4 | $\alpha/\beta(1 \rightarrow 3)$ linkage bead |
| B | SP1r | C3, H3, O3, C4, H4, O4, HO4 | $\alpha/\beta(1 \rightarrow 4)$ linkage bead |
| B | SN6 | C4, H4, O4, HO4, C5, H5 | Neu5Ac ring bead |
| C | SP1r | O5, C5, H5, C6, H61, H62, O6, HO6 | Standard pyranose bead |
| C | SN6r | O5, C5, H5, C6, H61, H62, O6 | $\alpha/\beta(1 \rightarrow 6)$ linkage bead |
| C | SP1r | O6, C6, H6, C7, H7, O7, HO7 | Neu5Ac bead |
| C | SN4ar | O5, C5, H5, C6, H61, H62, H63 | Fucose bead |
| D | SP3d | N, HN, C, O, CT, HT1, HT2, HT3 | N-acetyl (NHAc) group |
| E | SP3 | C8, H8, O8, HO8, C9, H91, H92, O9, HO9 | Neu5Ac terminal bead |

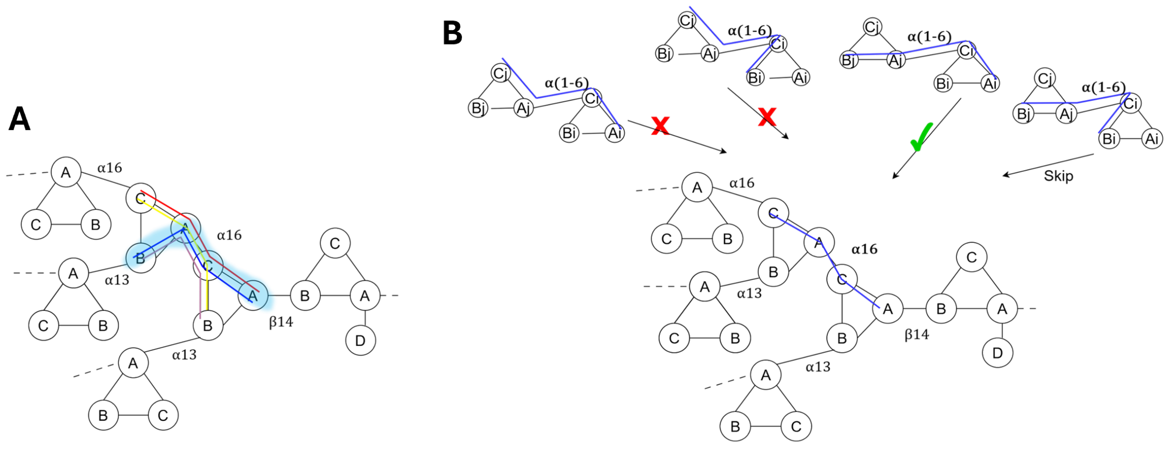


Figure S2: **A.** Two dihedral definitions per linkage unit in the branch points can overlap with other two dihedrals, as demonstrated in $\alpha(1-6)$ definition. **B.** A brute-force screening of dihedral configurations was shown for the $\alpha(1-6)$ linkage unit in the branch region. The tested ones leading to instabilities were rejected (red X), whereas the first configuration that produced stable simulations across all glycans was accepted (green tick). The same strategy was adopted for all linkage types, resulting in final configurations in Table 1.

Table S2: Generic bonded parameters for key glycosidic linkages in N-glycans

| Linkage | Bond | Angles | Dihedral |
|---|---|---|---|
| Neu5Ac($\alpha$2–3)Gal | B'–A: 0.314 / 8155 | A'–B'–A: 103 / 137<br>B'–A–B: 99 / 294 | A'–B'–A–B: -148 / 54 |
| Neu5Ac($\alpha$2–6)Gal | C'–A: 0.334 / 10085 | A'–C'–A: 149 / 192<br>C'–A–B: 135 / 150 | A'–C'–A–B: 125 / 8 |
| Fuc($\alpha$1–6)GlcNAc | C–A: 0.376 / 5169 | A'–C'–A: 131 / 147<br>C'–A–B: 102 / 220 | A'–C'–A–B: -93 / 12 |
| Gal($\beta$1–4)GlcNAc | B'–A: 0.358 / 5607 | A'–B'–A: 141 / 276<br>B'–A–C: 68 / 298 | A'–B'–A–C: -141 / -26 |
| GlcNAc($\beta$1–4)GlcNAc | B'–A: 0.309 / 17066 | A'–B'–A: 150 / 314<br>B'–A–C: 79 / 320 | A'–B'–A–C: 31 / 8 |
| Man($\beta$1–4)GlcNAc | B'–A: 0.344 / 9416 | A'–B'–A: 147 / 238<br>B'–A–C: 72 / 313 | A'–B'–A–C: -150 / -19 |
| Man($\alpha$1–2)Man | A'–A: 0.359 / 5379 | B'–A'–A: 98 / 54<br>A'–A–C: 59 / 274 | B'–A'–A–C: 38 / 7 |
| Man($\alpha$1–3)Man | B'–A: 0.336 / 4729 | A'–B'–A: 107 / 143<br>B'–A–B: 115 / 130 | A'–B'–A–B: -143 / 15 |
| Man($\alpha$1–6)Man | C'–A: 0.374 / 4350 | A'–C'–A: 116 / 203<br>C'–A–B: 101 / 107 | A'–C'–A–B: 110 / 32 |
| GlcNAc($\beta$1–2)Man | A'–A: 0.286 / 17365 | C'–A'–A: 113 / 95<br>A'–A–C: 98 / 64 | C'–A'–A–C: -27 / 6 |
| GlcNAc($\beta$1–4)Man | B'–A: 0.310 / 15635 | A'–B'–A: 154 / 243<br>B'–A–C: 80 / 308 | A'–B'–A–C: 58 / 11 |
| GlcNAc($\beta$1–6)Man | C'–A: 0.337 / 28436 | A'–C'–A: 119 / 112<br>C'–A–C: 95 / 240 | A'–C'–A–C: -44 / 10 |

Each entry is reported as equilibrium value/force constant. Bond values are reported in nm, whereas angle and dihedral values are reported in degrees.

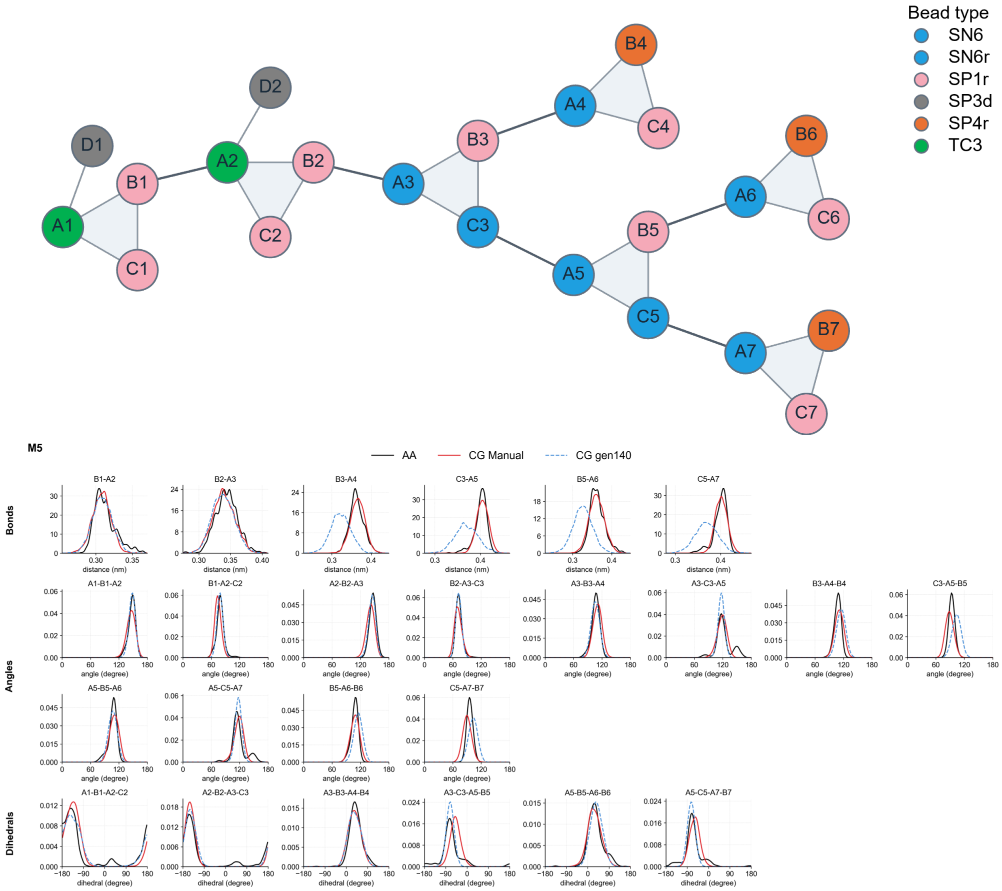


Figure S3: Coarse-grained representation of M5 glycan (top) with the color-coded beads indicating the bead types. The distributions of bonded interaction parameters (bottom) of generic CG gen140 models (CG Generic) are compared against the reference all-atom/pseudo-CG (AA) and the manually optimized models (CG Manual)

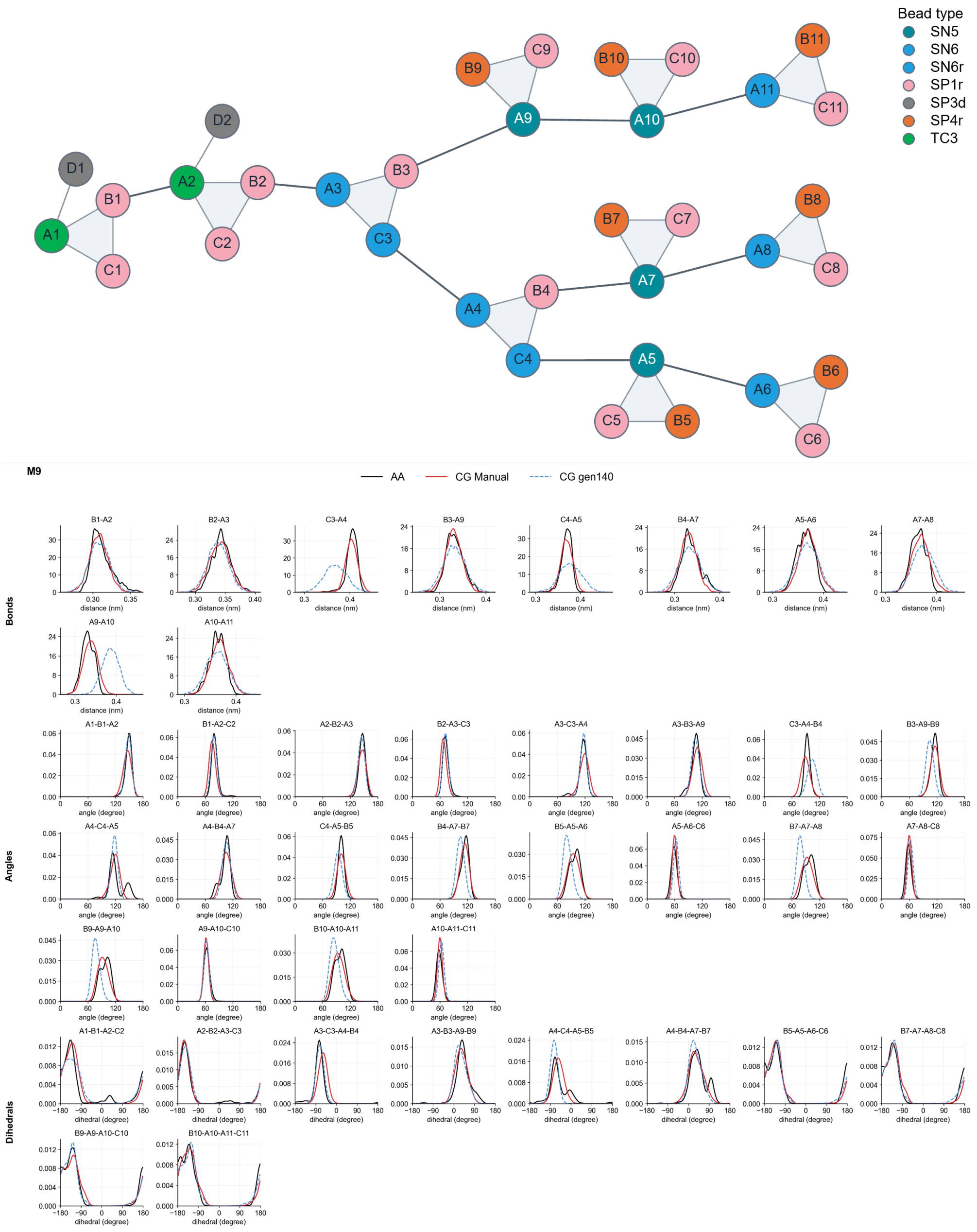


Figure S4: Coarse-grained representation of M9 glycan (top) with the color-coded beads indicating the bead types. The distributions of bonded interaction parameters (bottom) of generic CG gen140 models (CG Generic) are compared against the reference all-atom/pseudo-CG (AA) and the manually optimized models (CG Manual)

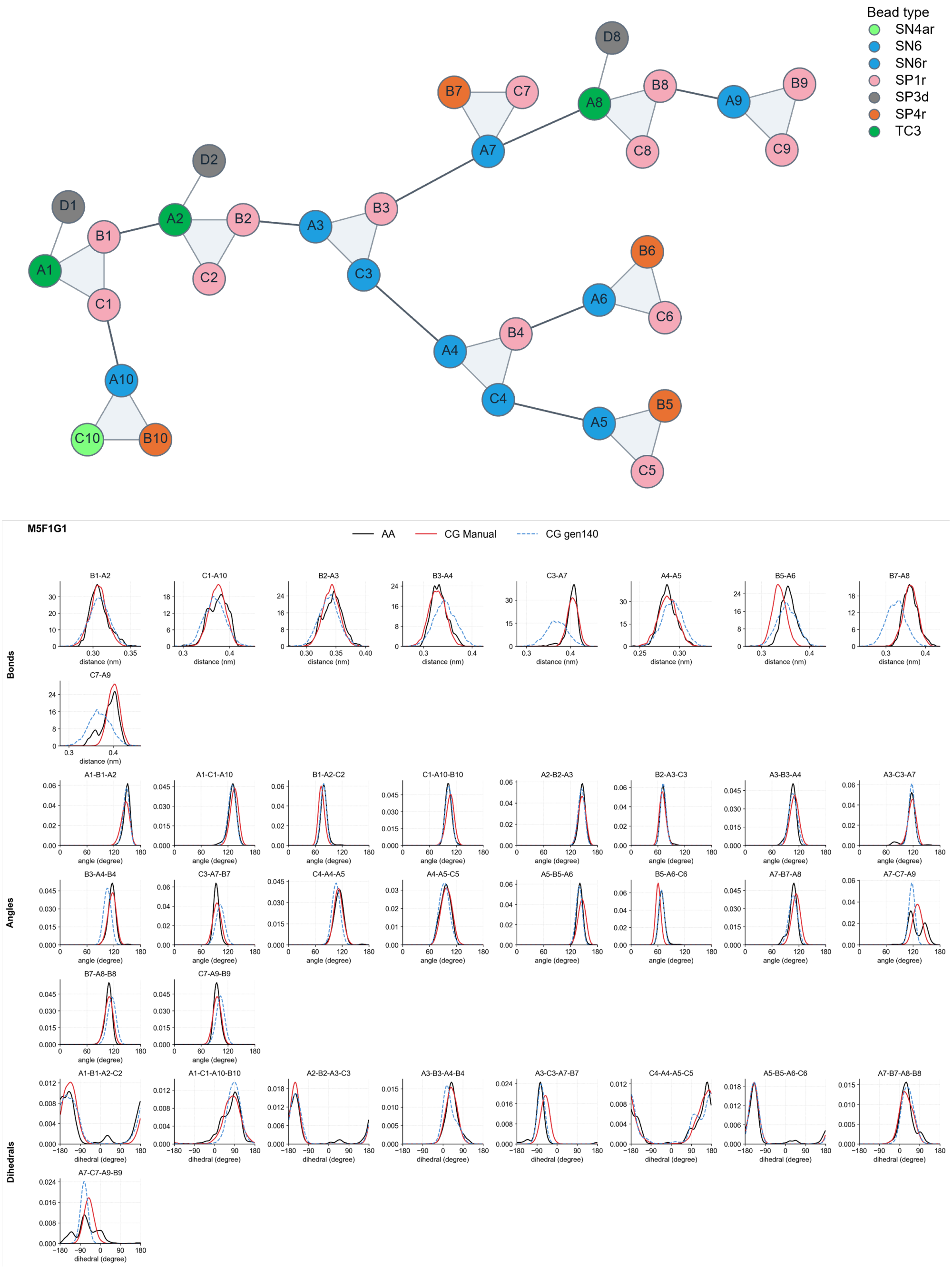


Figure S5: Coarse-grained representation of M5G1F1 glycan (top) with the color-coded beads indicating the bead types. The distributions of bonded interaction parameters (bottom) of generic CG gen140 models (CG Generic) are compared against the reference all-atom/pseudo-CG (AA) and the manually optimized models (CG Manual)

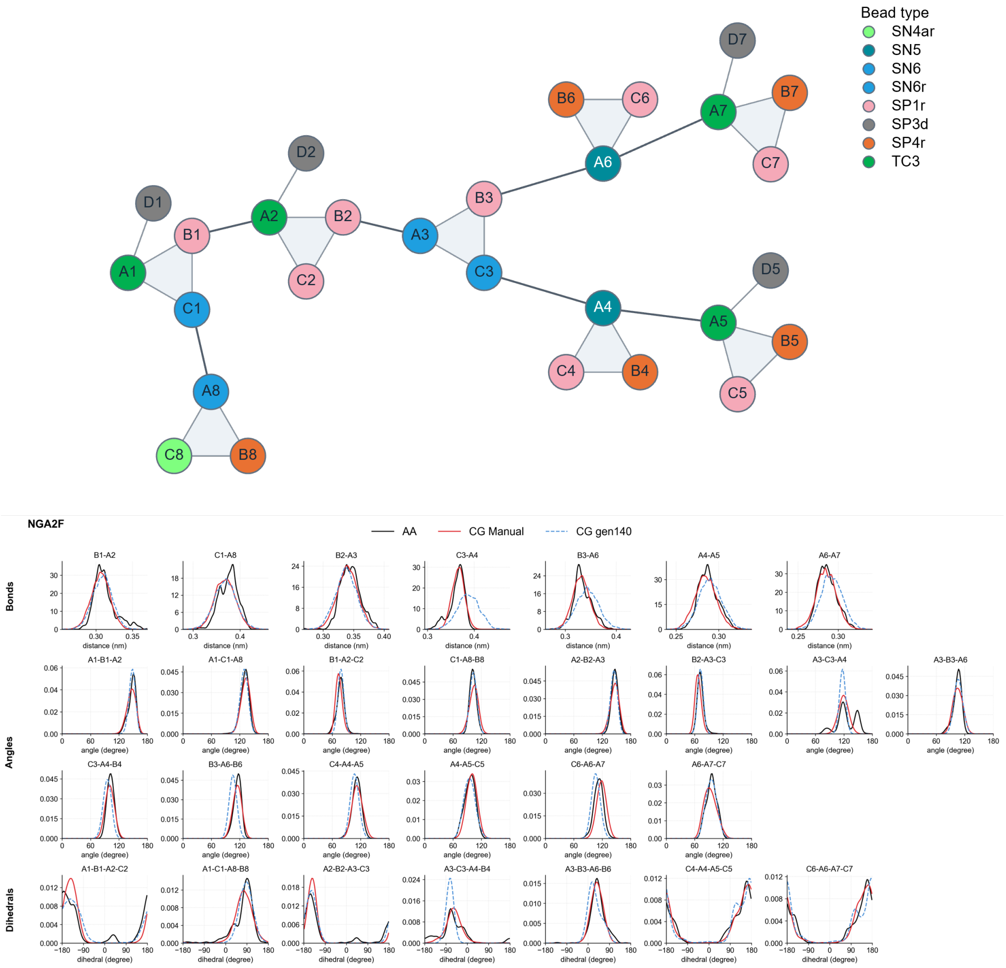


Figure S6: Coarse-grained representation of NGA2F glycan (top) with the color-coded beads indicating the bead types. The distributions of bonded interaction parameters (bottom) of generic CG gen140 models (CG Generic) are compared against the reference all-atom/pseudo-CG (AA) and the manually optimized models (CG Manual)

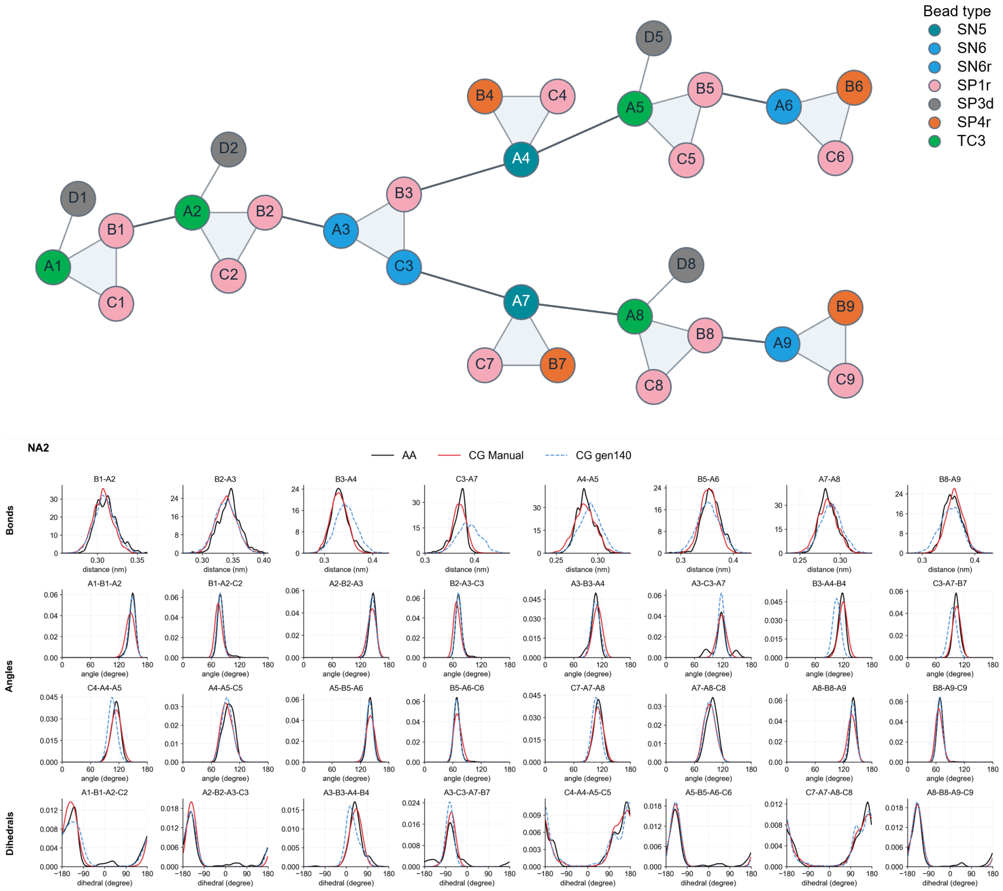


Figure S7: Coarse-grained representation of NA2 glycan (top) with the color-coded beads indicating the bead types. The distributions of bonded interaction parameters (bottom) of generic CG gen140 models (CG Generic) are compared against the reference all-atom/pseudo-CG (AA) and the manually optimized models (CG Manual)

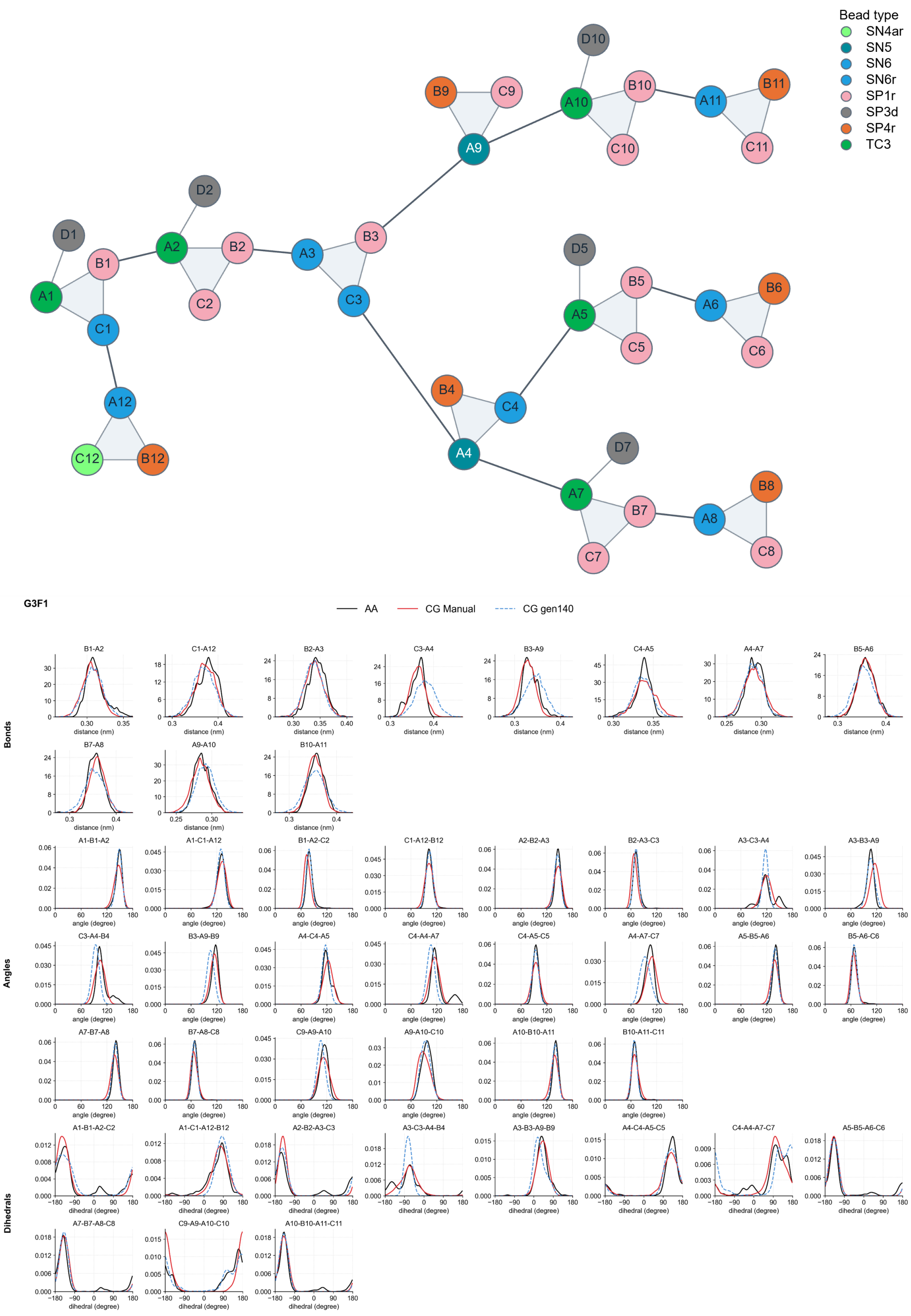


Figure S8: Coarse-grained representation of G3F1 glycan (top) with the color-coded beads indicating the bead types. The distributions of bonded interaction parameters (bottom) of generic CG gen140 models (CG Generic) are compared against the reference all-atom/pseudo-CG (AA) and the manually optimized models (CG Manual)

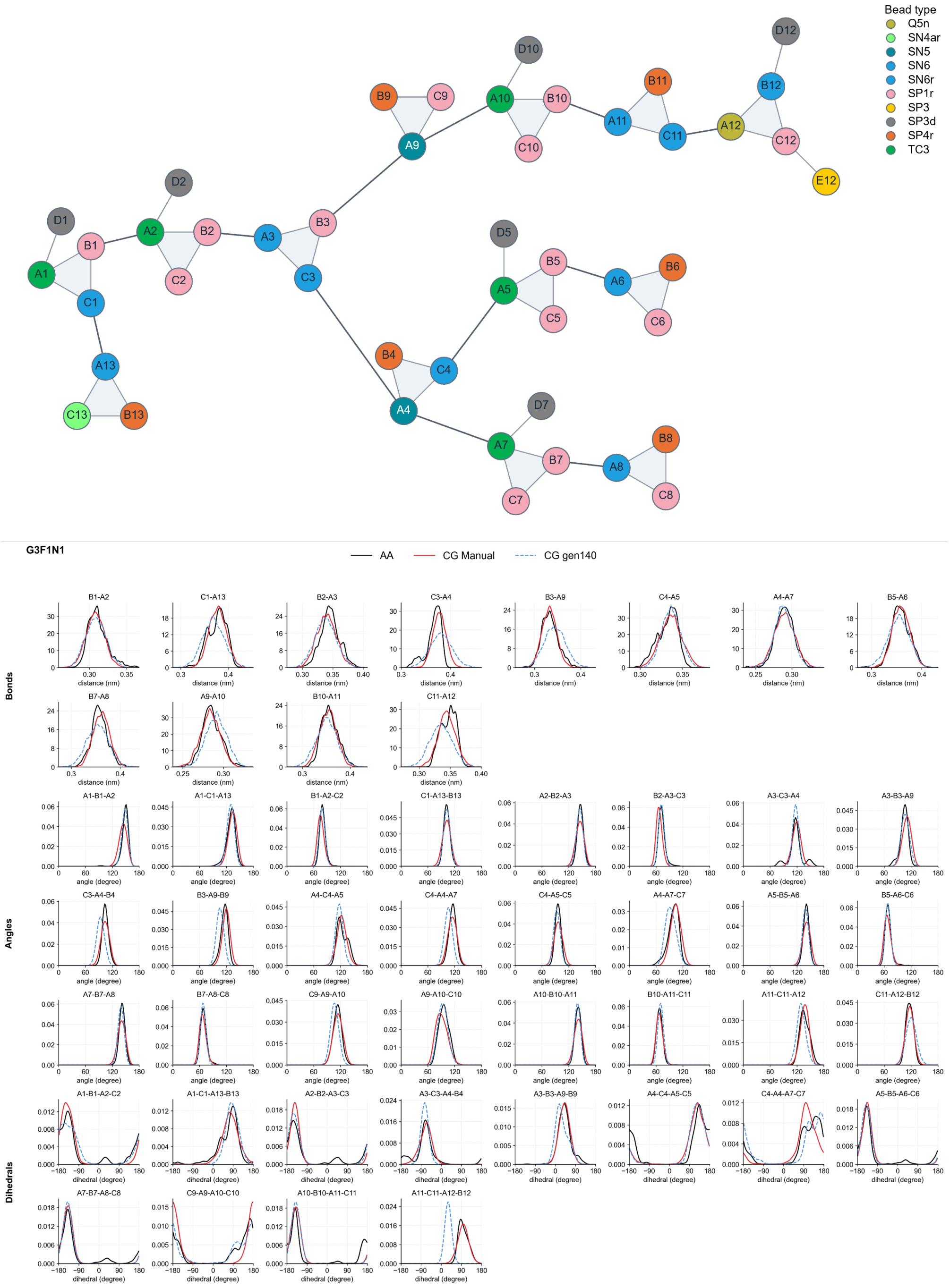

Figure S9: Coarse-grained representation of G3F1N1 glycan (top) with the color-coded beads indicating the bead types. The distributions of bonded interaction parameters (bottom) of generic CG gen140 models (CG Generic) are compared against the reference all-atom/pseudo-CG (AA) and the manually optimized models (CG Manual)

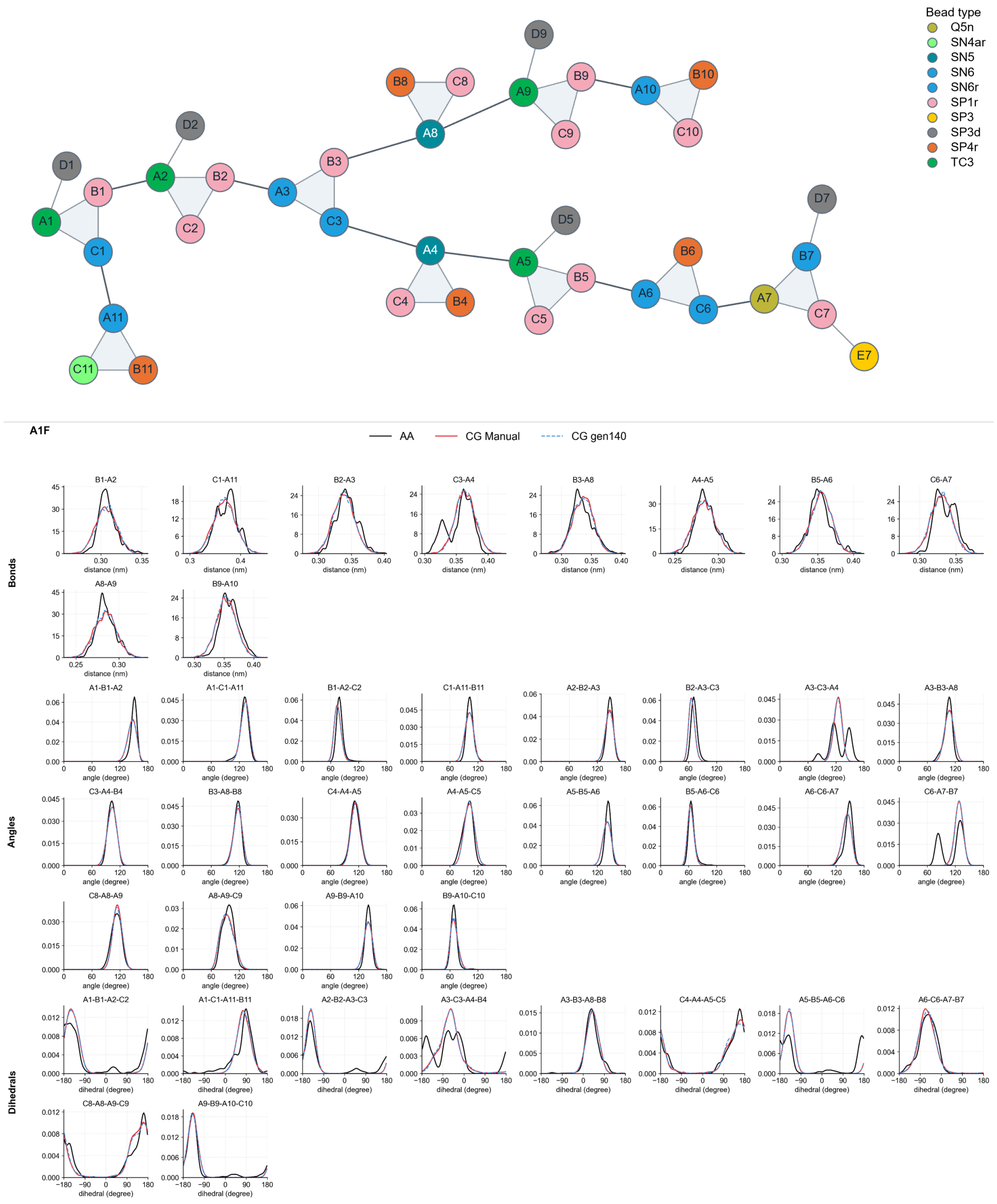


Figure S10: Coarse-grained representation of A1F glycan (top) with the color-coded beads indicating the bead types. The distributions of bonded interaction parameters (bottom) of generic CG gen140 models (CG Generic) are compared against the reference all-atom/pseudo-CG (AA) and the manually optimized models (CG Manual)

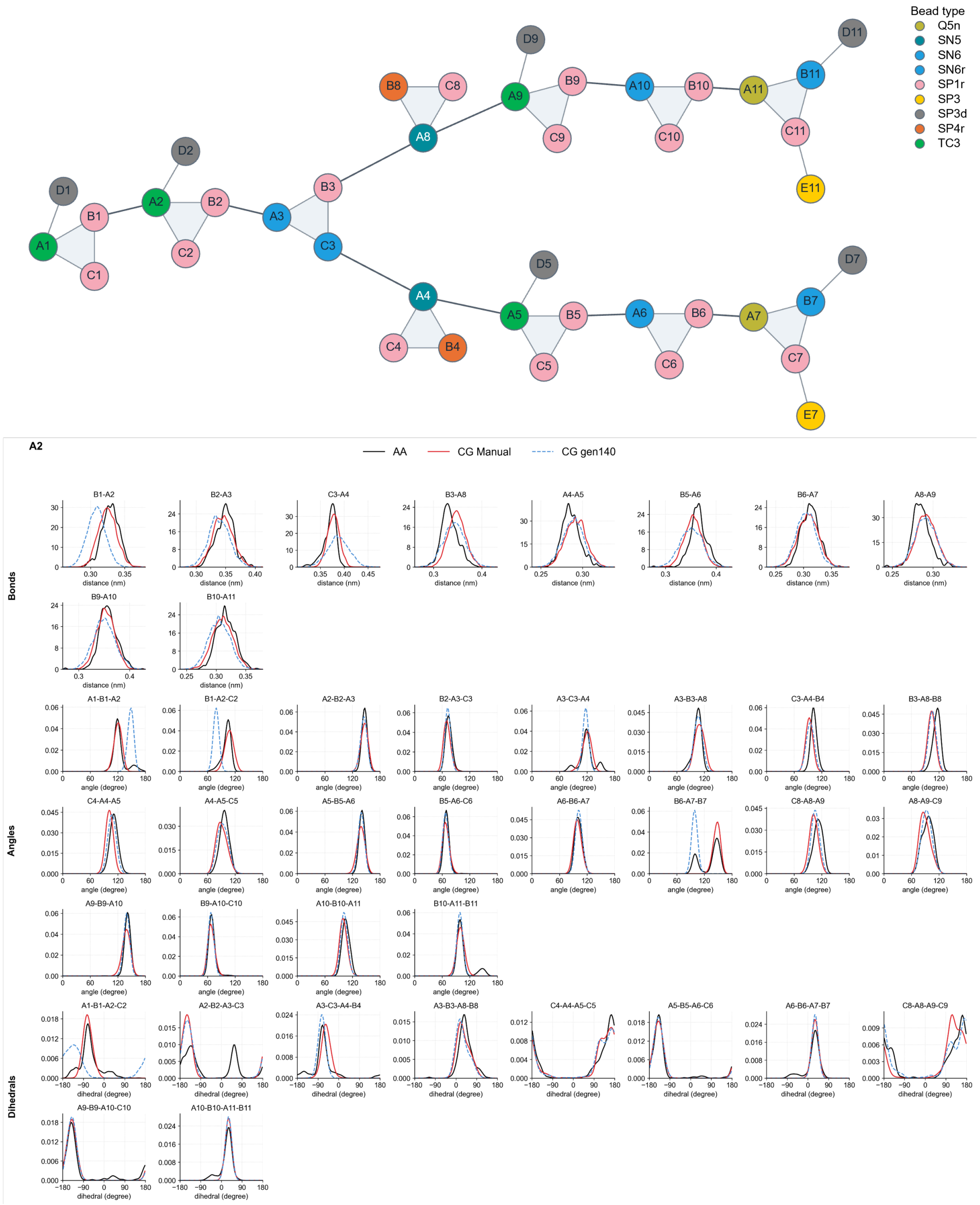


Figure S11: Coarse-grained representation of A2 glycan (top) with the color-coded beads indicating the bead types. The distributions of bonded interaction parameters (bottom) of generic CG gen140 models (CG Generic) are compared against the reference all-atom/pseudo-CG (AA) and the manually optimized models (CG Manual)

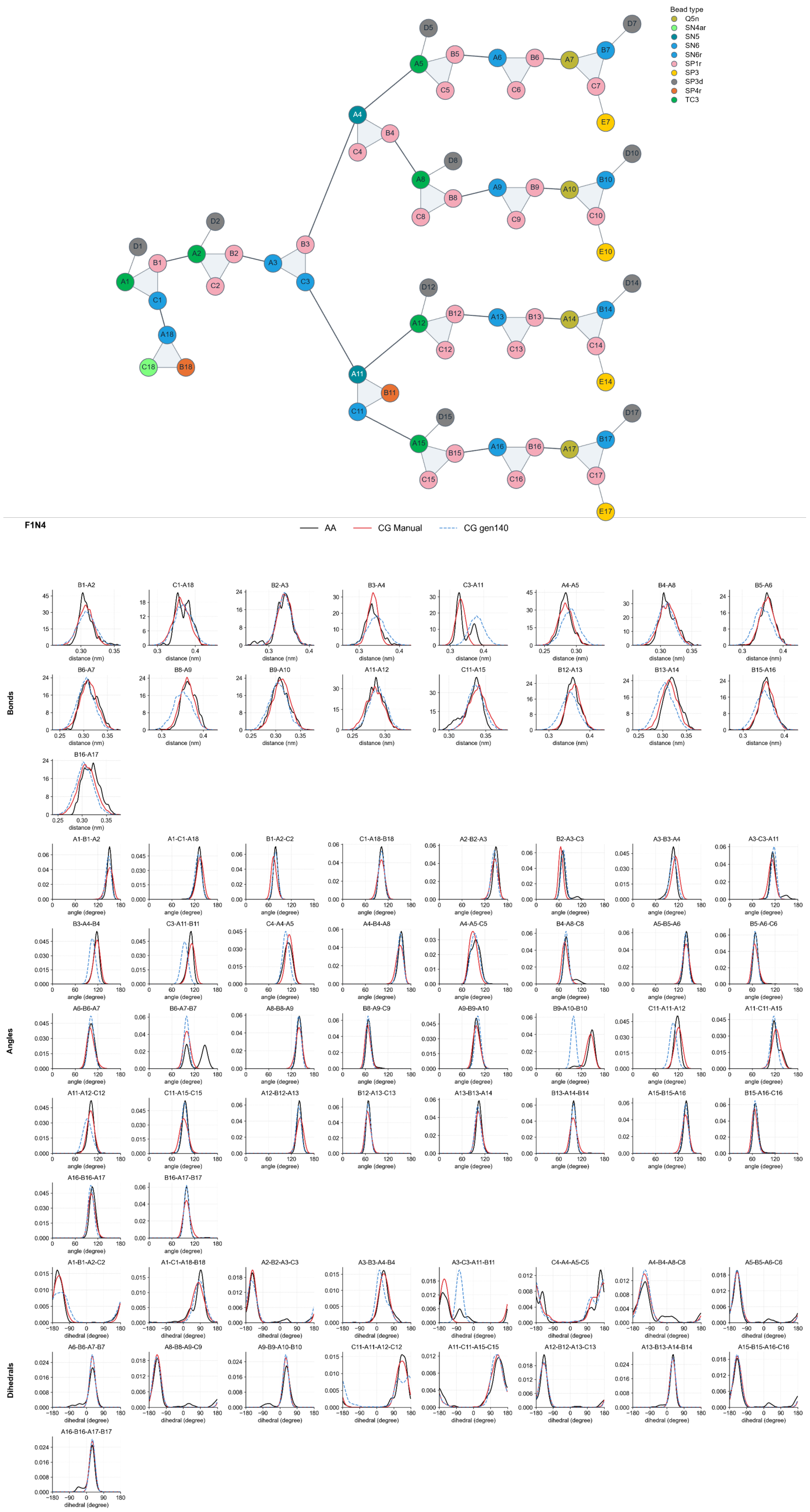


Figure S12: Coarse-grained representation of F1N4 glycan (top) with the color-coded beads indicating the bead types. The distributions of bonded interaction parameters (bottom) of generic CG gen140 models (CG Generic) are compared against the reference all-atom/pseudo-CG (AA) and the manually optimized models (CG Manual)

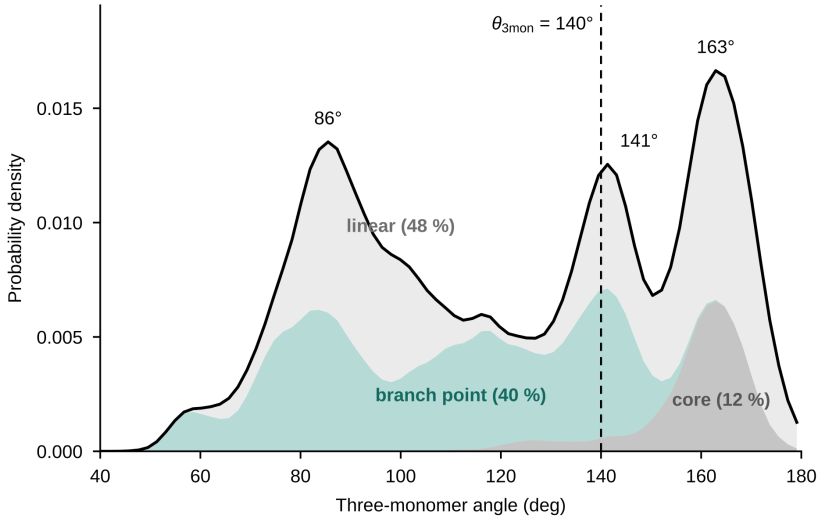


Figure S13: Pooled three-monomer angle distribution of the mapped AA trajectories (as in Fig. 4B), decomposed by the position of the central monomer: linear segments (48% of samples), branch points (40%) and the chitobiose core (12%). The dashed line marks $\theta_{3\mathrm{mon}} = 140°$.

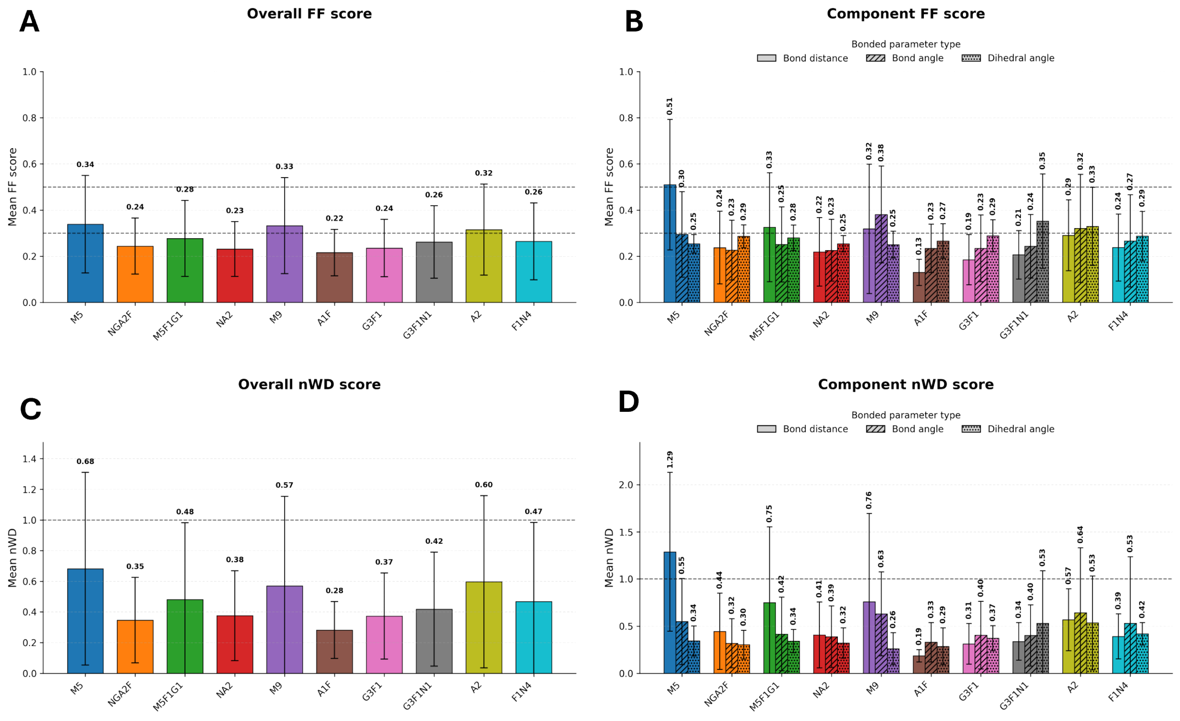


Figure S14: Comparison of CG gen140 and AA bonded parameter distributions for the ten glycans using FF and nWD metrics. **A** Overall FF scores. **B** FF score components. **C** Overall nWD scores. **D** nWD score components.

Table S3: Comparison of solvent-accessible surface area between AA, CG genNo, and CG gen140

| Glycan | AA | CG genNo | $\Delta_{\text{genNo}}$ (%) | CG gen140 | $\Delta_{\text{gen140}}$ (%) |
|---|---|---|---|---|---|
| M5 | 15.87 ± 0.60 | 15.19 ± 0.92 | -4.3 | 16.64 ± 0.80 | 4.9 |
| NGA2F | 18.27 ± 0.61 | 17.14 ± 1.08 | -6.2 | 18.64 ± 1.03 | 2.0 |
| M5F1G1 | 21.30 ± 0.70 | 19.86 ± 1.31 | -6.7 | 21.71 ± 1.32 | 1.9 |
| NA2 | 20.66 ± 0.70 | 18.91 ± 1.15 | -8.5 | 20.82 ± 1.11 | 0.8 |
| M9 | 21.29 ± 0.78 | 18.95 ± 1.14 | -11.0 | 22.19 ± 1.09 | 4.2 |
| A1F | 23.84 ± 1.60 | 22.65 ± 1.61 | -5.0 | 24.78 ± 1.66 | 3.9 |
| G3F1 | 25.36 ± 1.17 | 23.65 ± 1.60 | -6.7 | 25.95 ± 1.56 | 2.3 |
| G3F1N1 | 27.63 ± 1.29 | 25.71 ± 1.82 | -7.0 | 28.74 ± 1.93 | 4.0 |
| A2 | 26.07 ± 0.85 | 24.00 ± 1.52 | -7.9 | 25.79 ± 1.50 | -1.1 |
| F1N4 | 38.93 ± 2.38 | 35.28 ± 2.45 | -9.4 | 38.56 ± 2.50 | -0.9 |

Values are reported as mean ± standard deviation. $\Delta$ is calculated as $100 \times (\text{CG} - \text{AA})/\text{AA}$. SASA is reported in $\text{nm}^2$.

Table S4: Comparison of radius of gyration between AA, CG genNo, and CG gen140

| Glycan | AA | CG genNo | $\Delta_{\text{genNo}}$ (%) | CG gen140 | $\Delta_{\text{gen140}}$ (%) |
|---|---|---|---|---|---|
| M5 | 0.70 ± 0.03 | 0.70 ± 0.05 | -0.8 | 0.79 ± 0.05 | 12.2 |
| NGA2F | 0.78 ± 0.03 | 0.73 ± 0.05 | -5.8 | 0.82 ± 0.06 | 5.4 |
| M5F1G1 | 0.86 ± 0.03 | 0.80 ± 0.06 | -7.6 | 0.89 ± 0.06 | 2.8 |
| NA2 | 0.88 ± 0.04 | 0.78 ± 0.05 | -11.3 | 0.88 ± 0.05 | -0.4 |
| M9 | 0.81 ± 0.02 | 0.73 ± 0.04 | -9.3 | 0.88 ± 0.04 | 8.5 |
| A1F | 0.94 ± 0.07 | 0.87 ± 0.07 | -7.4 | 0.94 ± 0.07 | 0.1 |
| G3F1 | 0.94 ± 0.04 | 0.87 ± 0.05 | -7.5 | 0.95 ± 0.06 | 0.8 |
| G3F1N1 | 1.01 ± 0.04 | 0.92 ± 0.07 | -8.8 | 0.99 ± 0.07 | -1.8 |
| A2 | 1.07 ± 0.06 | 0.92 ± 0.08 | -13.4 | 1.01 ± 0.07 | -5.0 |
| F1N4 | 1.22 ± 0.05 | 1.07 ± 0.07 | -11.9 | 1.16 ± 0.07 | -4.8 |

Values are reported as mean ± standard deviation. $\Delta$ is calculated as $100 \times (\text{CG} - \text{AA})/\text{AA}$. $R_g$ is reported in nm.

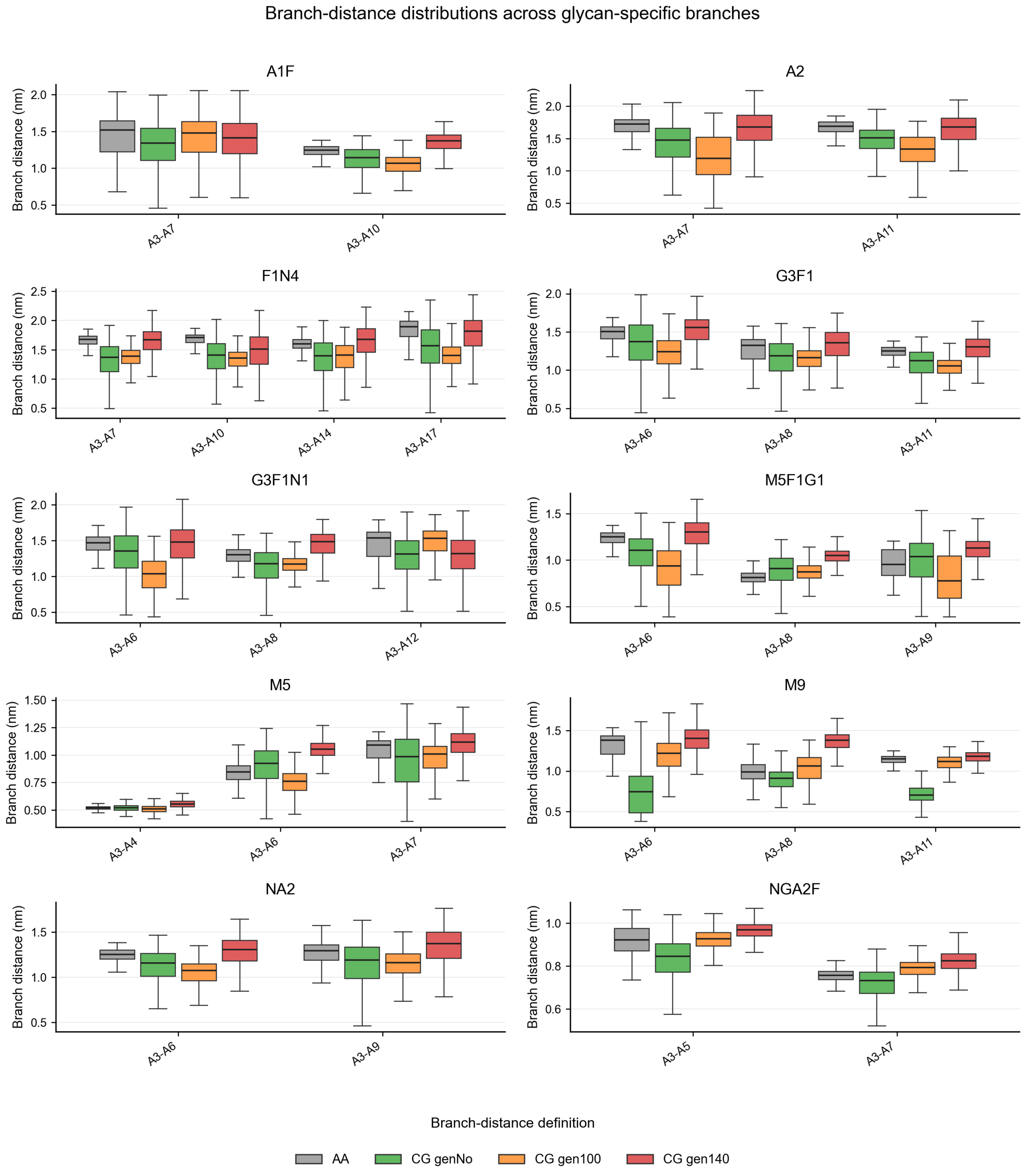


Figure S15: Branch distance distributions for the ten glycans represented as boxplots, comparing the three generic CG models (CG genNo, CG gen100, and CG gen140) with the AA reference

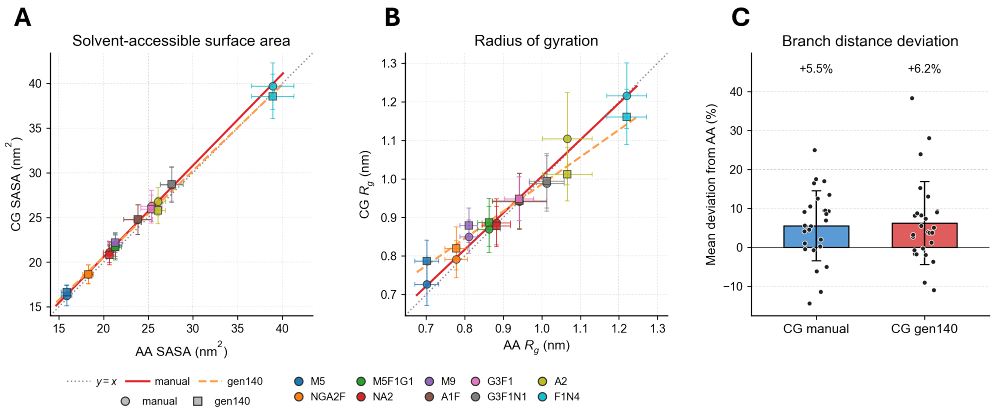


Figure S16: Assessment of manually CG-optimized GMR (*CG manual*) parametrization across the ten N-glycans relative to the AA references. **A.** Parity plot comparing SASA. **B.** The parity plot comparing averaged $R_g$ across all glycans. (C.) Absolute mean branch- distance difference from the AA reference for the *CG manual* and *gen140* models. The mean deviations against AA modesl were reported above the bars with the grey shaded region indicates deviations within ±5%

Table S5: Comparison of CG Manual SASA and radius of gyration relative to AA and CG gen140

| Glycan | SASA | | | $R_g$ | | |
|---|---|---|---|---|---|---|
| | CG Manual | $\Delta_{\mathrm{AA}}$ (%) | $\Delta_{\mathrm{Gen140}}$ (%) | CG Manual | $\Delta_{\mathrm{AA}}$ (%) | $\Delta_{\mathrm{Gen140}}$ (%) |
| M5 | 16.24 ± 1.12 | 2.4 | -2.4 | 0.73 ± 0.06 | 3.5 | -7.7 |
| NGA2F | 18.64 ± 1.08 | 2.0 | 0.0 | 0.79 ± 0.05 | 1.8 | -3.4 |
| M5F1G1 | 21.71 ± 1.45 | 1.9 | 0.0 | 0.87 ± 0.06 | 0.7 | -2.0 |
| NA2 | 21.13 ± 1.19 | 2.3 | 1.5 | 0.89 ± 0.06 | 0.4 | 0.9 |
| M9 | 21.98 ± 1.33 | 3.2 | -1.0 | 0.85 ± 0.04 | 4.9 | -3.3 |
| A1F | 24.74 ± 1.67 | 3.8 | -0.1 | 0.94 ± 0.07 | -0.1 | -0.2 |
| G3F1 | 26.30 ± 1.73 | 3.7 | 1.4 | 0.95 ± 0.06 | 0.8 | -0.0 |
| G3F1N1 | 28.64 ± 1.97 | 3.7 | -0.3 | 0.99 ± 0.07 | -2.4 | -0.6 |
| A2 | 26.80 ± 1.57 | 2.8 | 3.9 | 1.10 ± 0.12 | 3.5 | 9.0 |
| F1N4 | 39.71 ± 2.60 | 2.0 | 3.0 | 1.22 ± 0.09 | -0.4 | 4.7 |

Values are reported as mean ± standard deviation for the CG manual model only. $\Delta_{\mathrm{AA}}$ is calculated as $100 \times (\text{CG Manual} - \text{AA})/\text{AA}$, and $\Delta_{\mathrm{Gen140}}$ as $100 \times (\text{CG Manual} - \text{CG Gen140})/\text{CG Gen140}$. SASA is reported in $\mathrm{nm}^2$ and $R_g$ in nm.

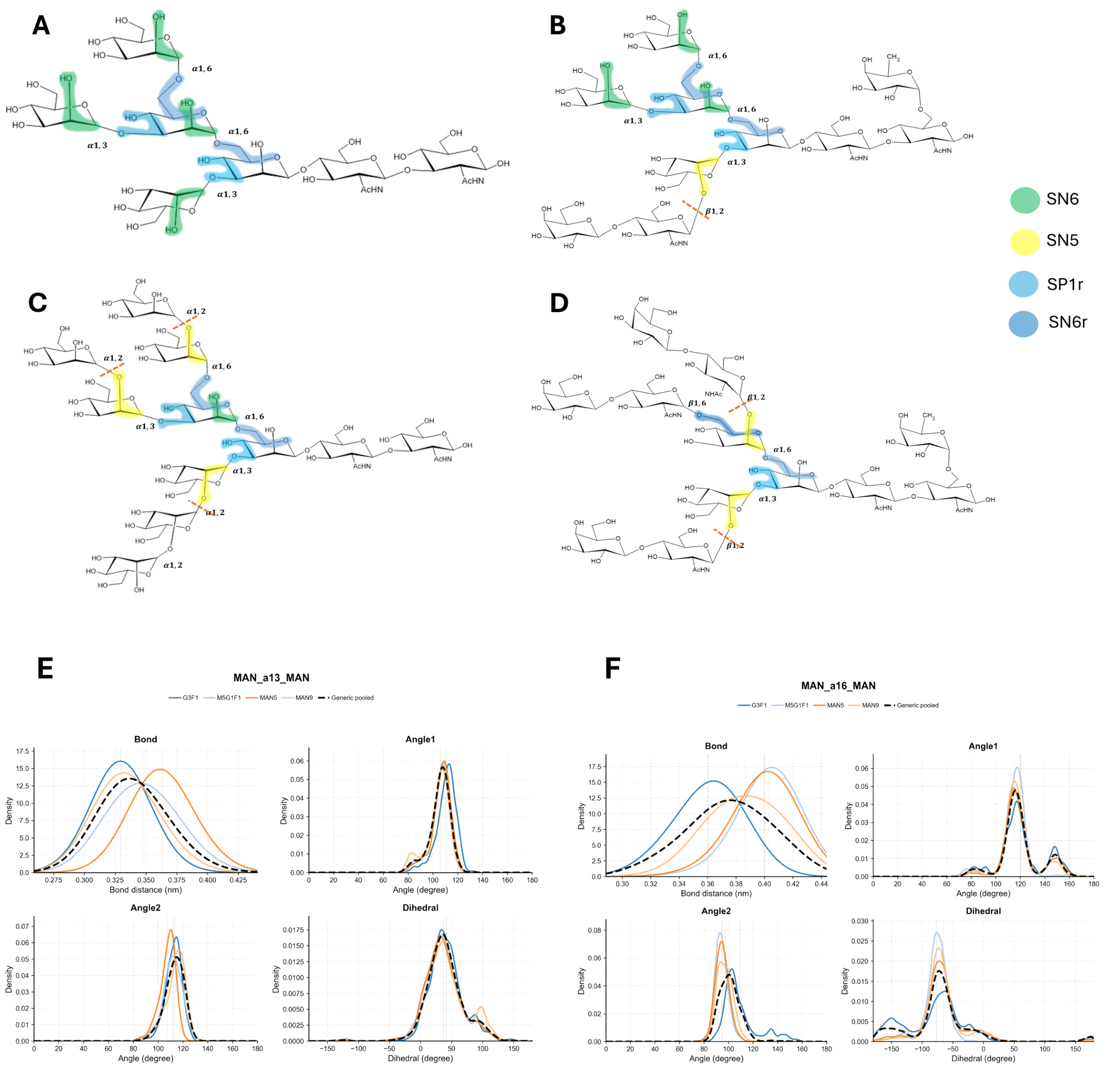


Figure S17: A difference in the mapping scheme of Man($\alpha1-3$)Man and Man($\alpha1-6$)Man linkages between M5 (**A**), as the smallest high-mannose glycan, with larger glycans: M9, M5G1F1, and G3F1, shown in **B**, **C**, and **D**, respectively. The red dashed lines in the three larger glycans show how the mapping is modified, depending on whether the mannose(s) is linked to the next monomer. The mapping choice largely contributes to a marginal discrepancy in the bond-length distributions of M5 and M5G1F1 in the generic parametrization (**E** and **F**), where the dashed black lines represent the pooled generic results. The generic parameters were inferred from the consensus distributions of 7 complex glycans against the 3 high-mannose glycans.

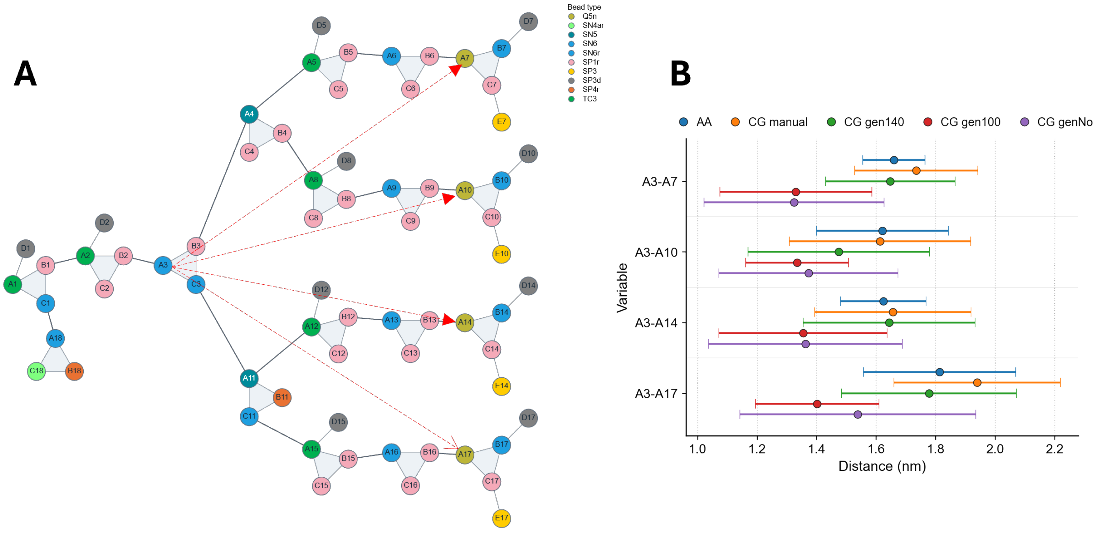


Figure S18: The definition of four branch distances in F1N4 glycan, which are shown as dashed red lines in **A**. The branch distance distributions for the three CG generic models (CG genNo, CG gen100, and CG gen140) are compared against the manually optimized models (CG Manual) and the atomistic reference distributions in **B**.

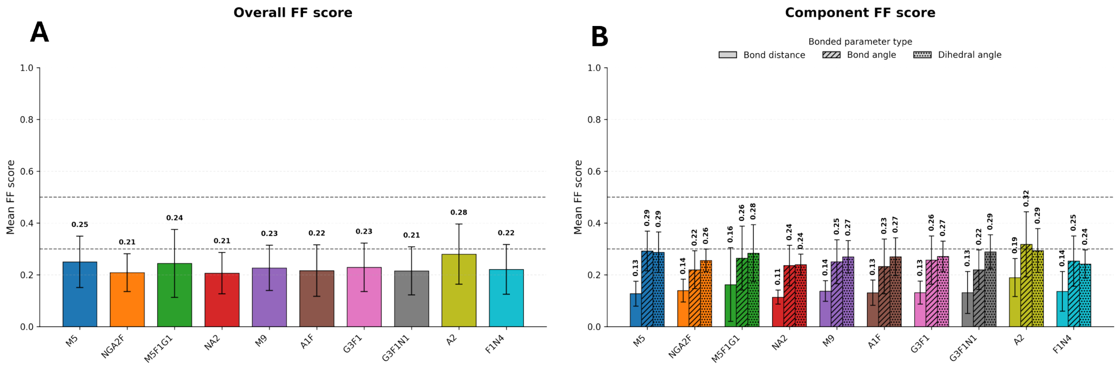


Figure S19: Comparison of the manually optimized CG models (CG Manual) and AA bonded parameter distributions for the ten glycans using FF scoring

Table S6: Generic bonded parameters for the Asn–GlcNAc linkage and neighboring aromatic amino acids

| Bonded variable | Potential | Parameters | Aromatic side-chain tuning |
|---|---|---|---|
| BB–SC1 | Harmonic bond | 0.264 / 18000 | – |
| SC1–A1 | Harmonic bond | 0.335 / 11000 | – |
| BB–SC1–A1 | Harmonic angle | 126 / 100 | – |
| SC1–A1–C1 | Harmonic angle | 100 / 100 | – |
| SC1–A1–B1 | ReB angle | 145 / 185 | – |
| −BB–BB–SC1–A1 | Periodic dihedral, $n = 1$ | 148 / 20 | – |
| +BB–BB–SC1–A1 | Periodic dihedral, $n = 1$ | −27 / 6 | – |
| BB–SC1–A1–C1 | Periodic dihedral, $n = 1$ | 5 / 12 | 160 / 20 |
| SC1–A1–B1–A2 | Periodic dihedral, $n = 1$ | 222 / 6 | 270 / 6 |

Each entry is reported as equilibrium value / force constant. Bond values are reported in nm, whereas angle and dihedral values are reported in degrees. Bonds were modeled using harmonic potentials. Angles were modeled using harmonic potentials, except for angles above 140°, which were modeled using restricted bending (ReB) potentials. All dihedrals were modeled using periodic potentials with multiplicity $n = 1$. We applied aromatic side-chain tuning only to the BB–SC1–A1–C1 and SC1–A1–B1–A2 dihedrals.

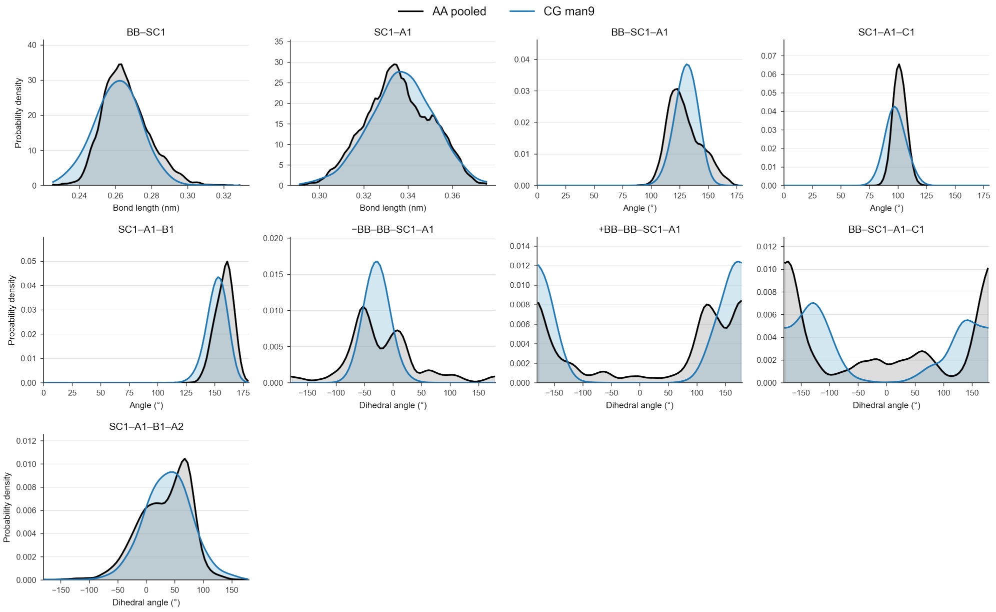


Figure S20: Comparison of the GlcNAc-Asn linkage parameter distributions obtained from simulations of tripeptide linked to M9 glycan between the mapped-AA (grey) and CG (blue).

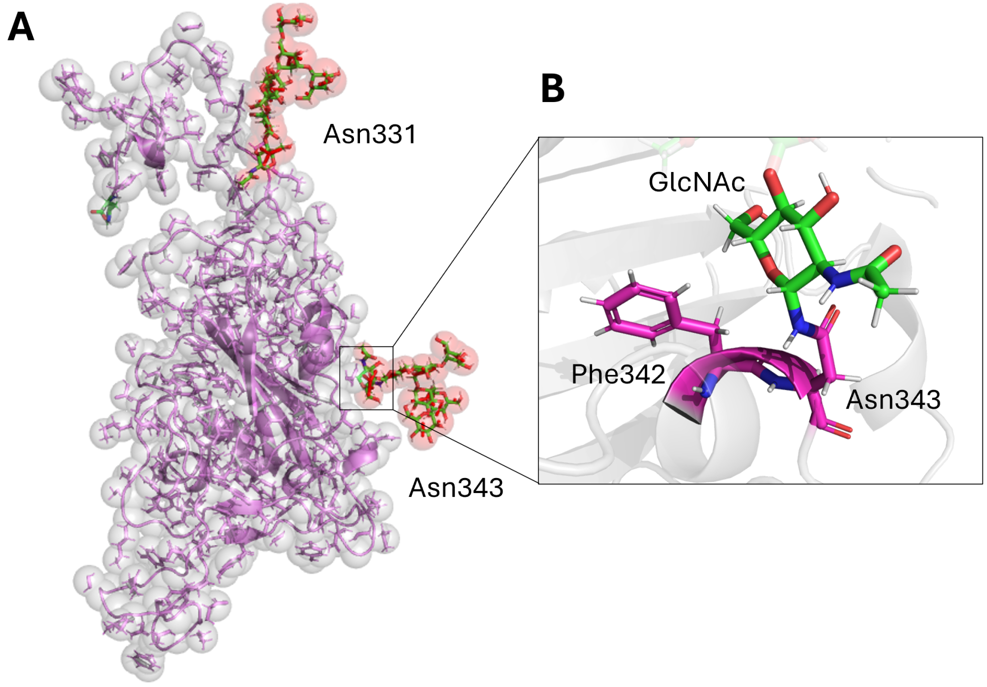


Figure S21: **A.** Atomistic (sticks) and CG (spheres) representations of the receptor-binding domain of SARS-Cov-2 spike protein used as the case study, where two glycosylation sites, Asn331 and Asn343, bearing M5 glycans. **B.** The suspected carbohydrate-aromatic interaction between the GlcNAc ring and the benzyl side chain of Phe342.

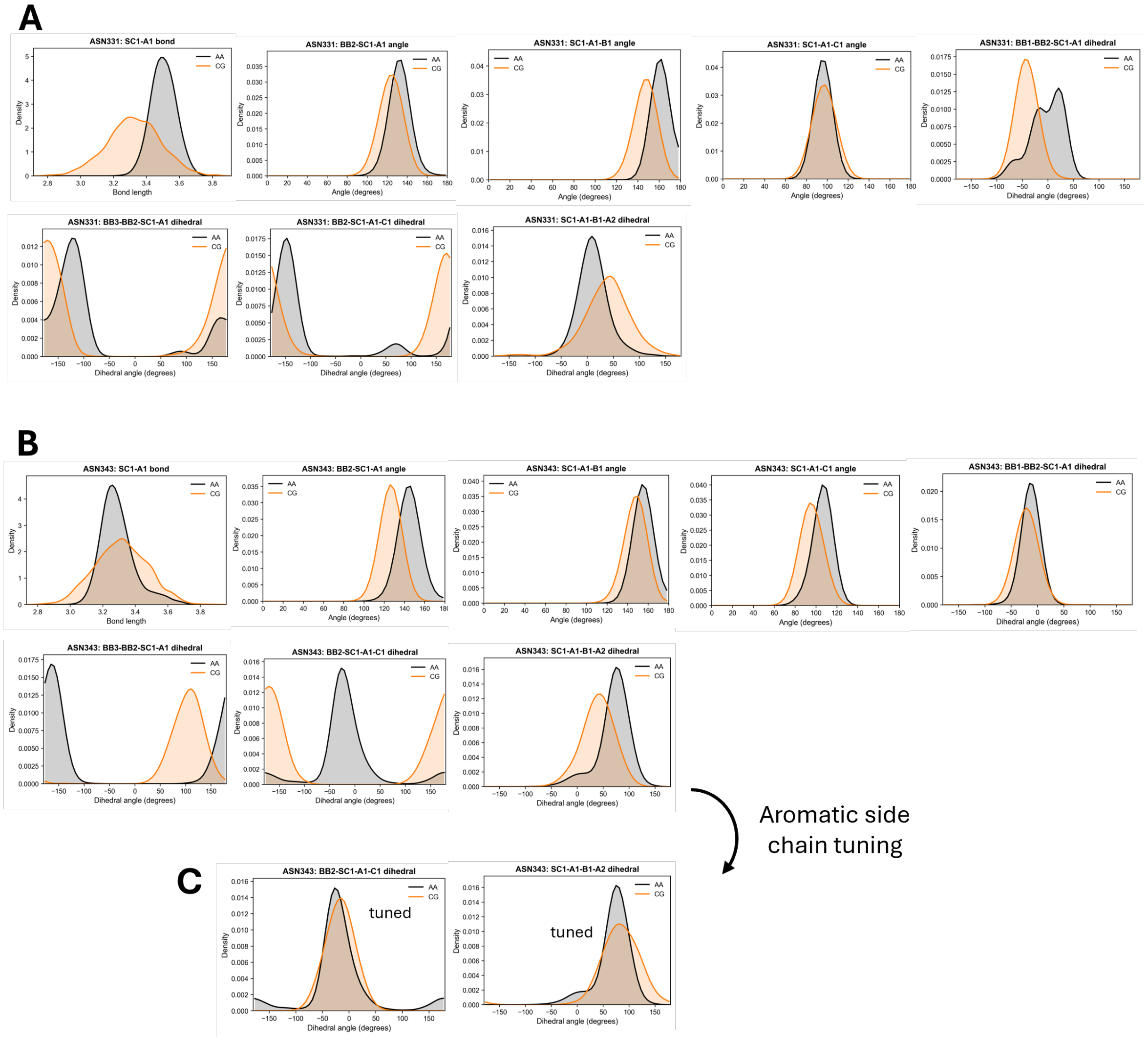


Figure S22: CG GlcNAc–Asn linkage parameter distributions extracted from the RBD+M5 system, focusing on the tripeptide segments of **A** Asn331 and **B** Asn343, shown in gray and compared with the corresponding AA reference distributions in orange. **C** Distributions of the two Asn343 dihedrals modified by aromatic side-chain tuning.

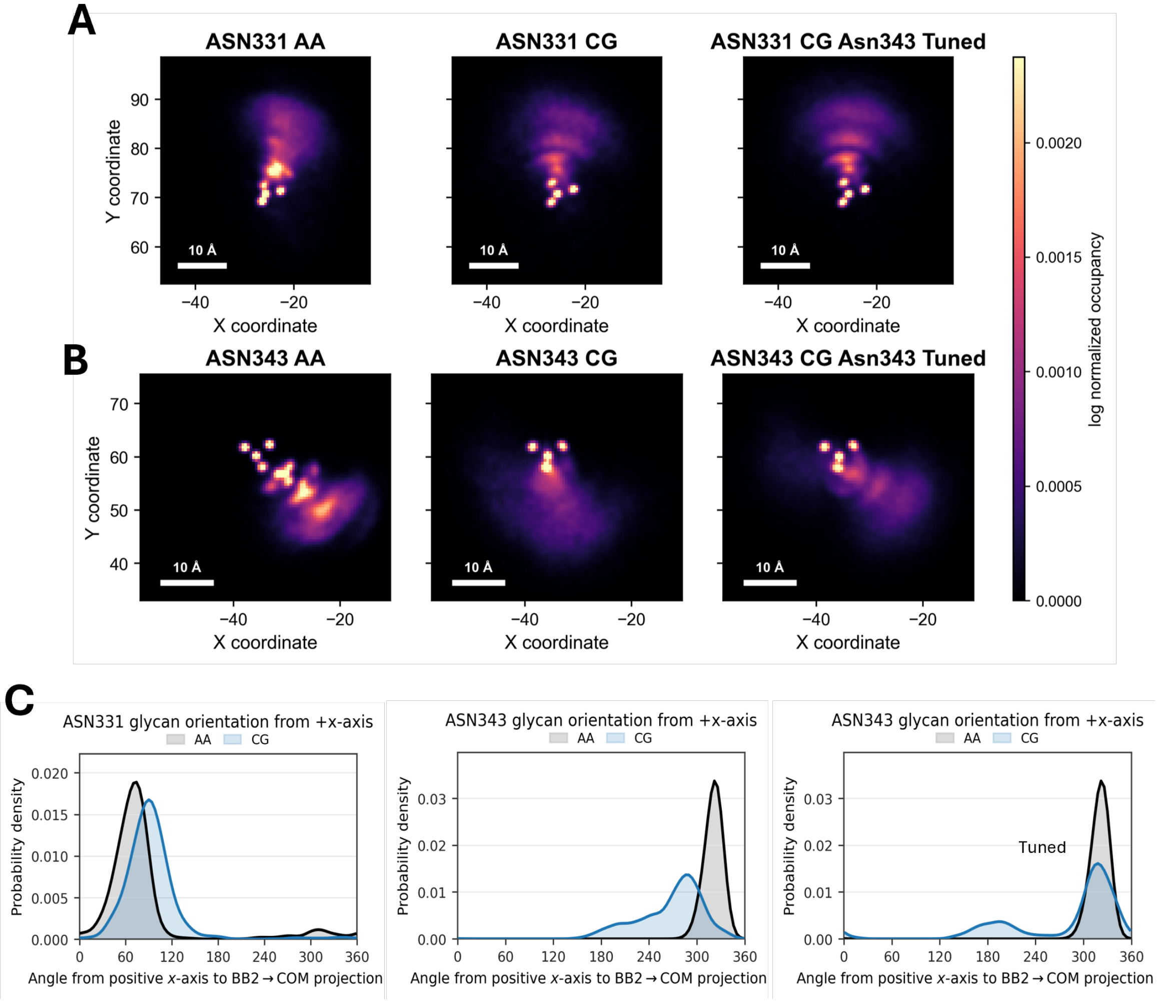


Figure S23: The $xy-$ projected occupancy maps of M5 glycans of the RBD protein attached to the two RBD glycosylation sites: **A** Asn331 and **B** Asn343 in the AA and CG models, including the CG model with tuned dihedrals at Asn343. **C** Glycan orientation distributions, calculated as the angle between the $+x$-axis and the vector from the glycosylation site to the glycan center of mass, for the AA, CG, and tuned CG models.

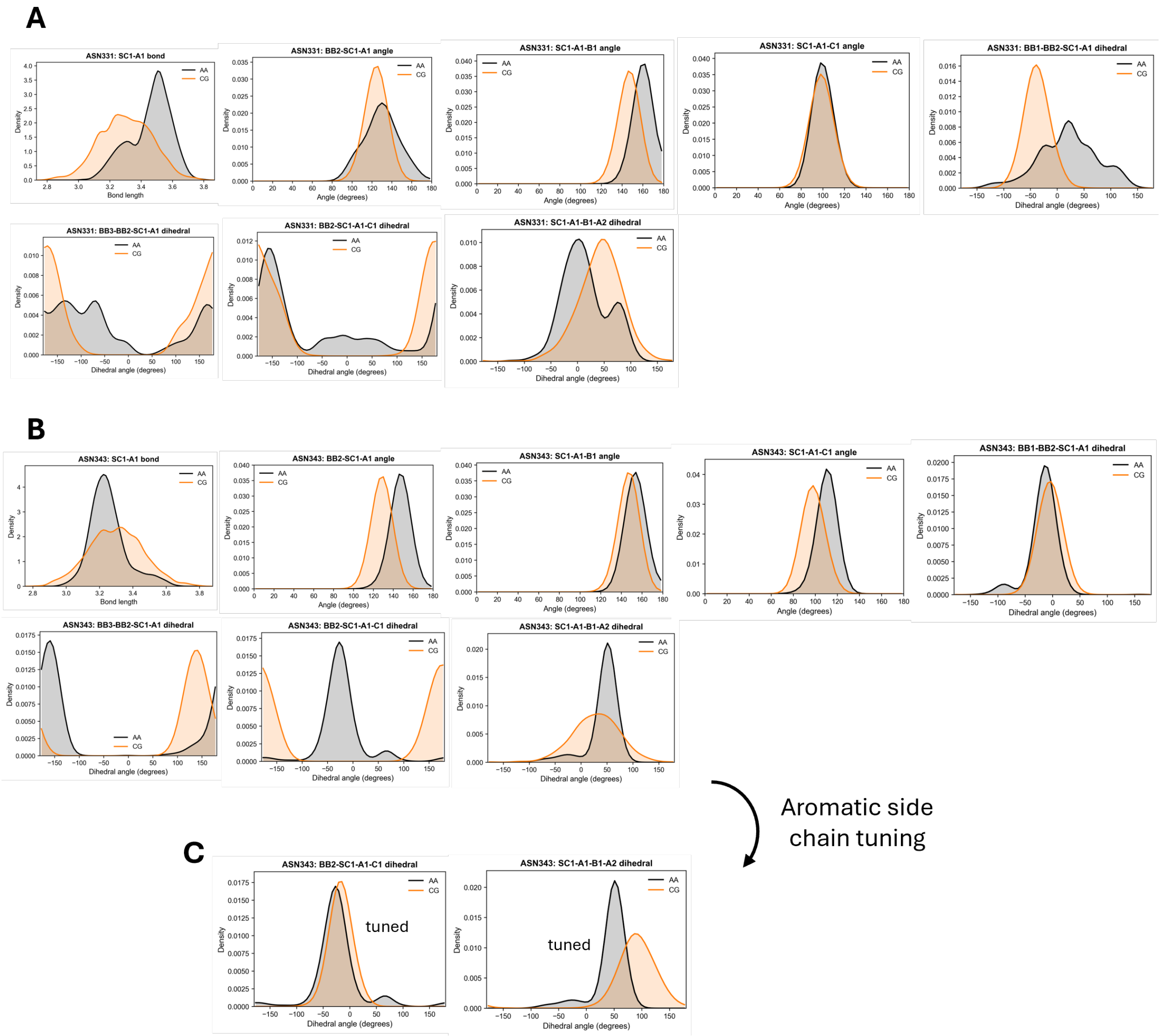


Figure S24: CG GlcNAc–Asn linkage parameter distributions extracted from the RBD+NGA2F system, focusing on the tripeptide segments of **A** Asn331 and **B** Asn343 are shown in gray and compared with the corresponding AA reference distributions in orange. **C** Distributions of the two Asn343 dihedrals tuned by aromatic neigboring amino acids

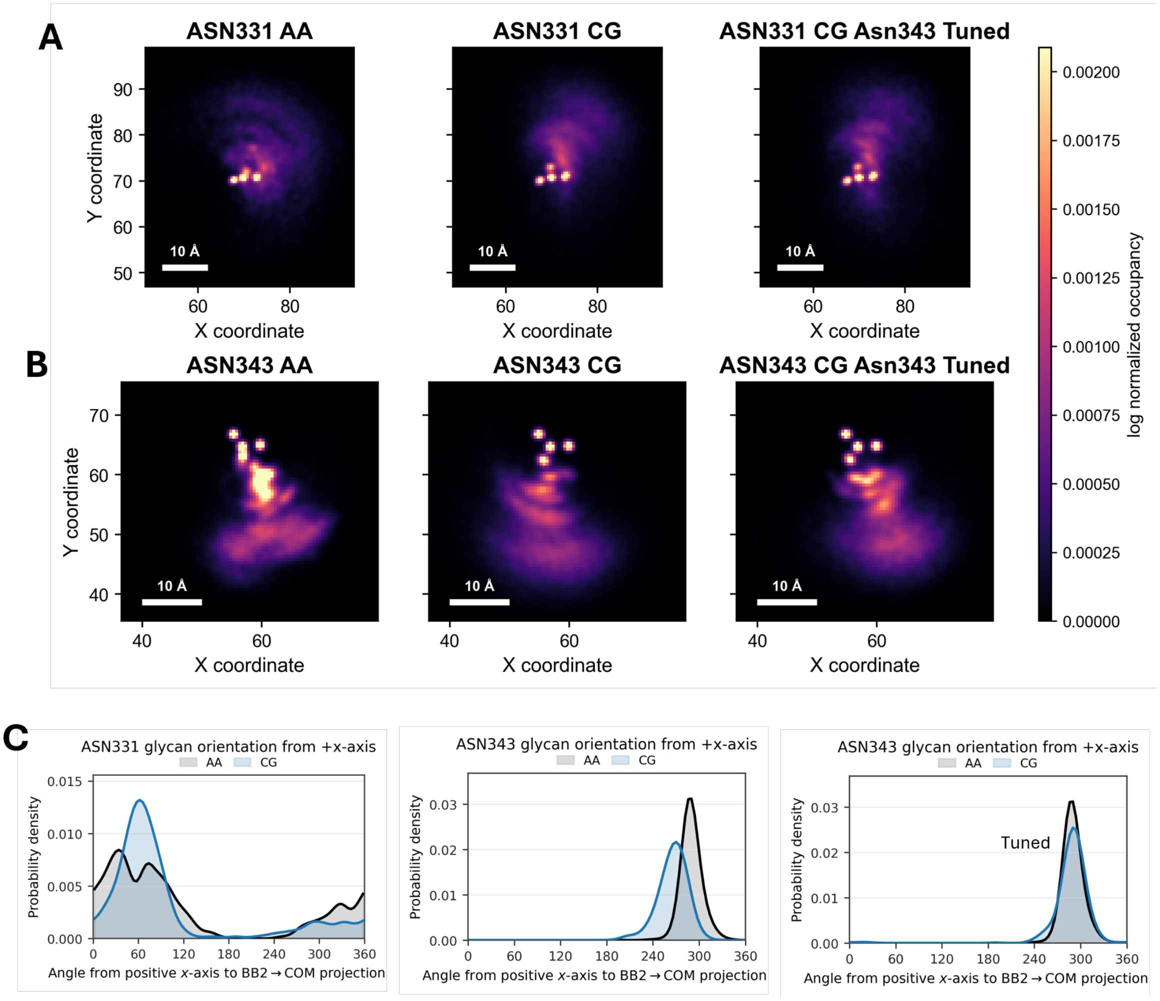


Figure S25: The $xy-$ projected occupancy maps of NGA2F glycans of the RBD protein attached to the two RBD glycosylation sites: **A** Asn331 and **B** Asn343 in the AA and CG models, including the CG model with tuned dihedrals at Asn343. **C** Glycan orientation distributions, calculated as the angle between the $+x$-axis and the vector from the glycosylation site to the glycan center of mass, for the AA, CG, and tuned CG models.

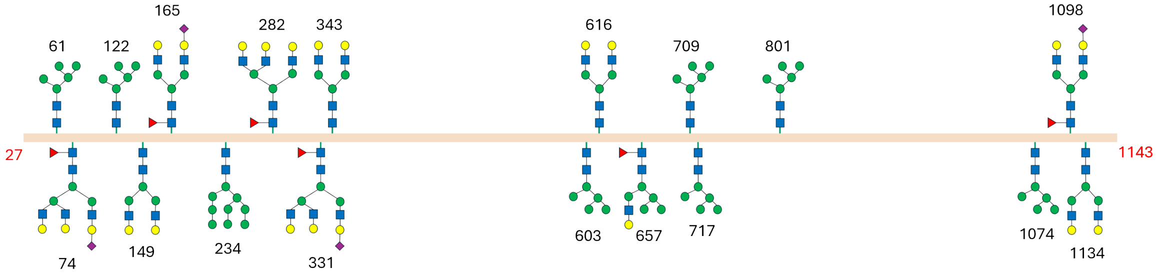


Figure S26: Glycan composition per subunit in SARS-Cov-2 Spike protein used in this work

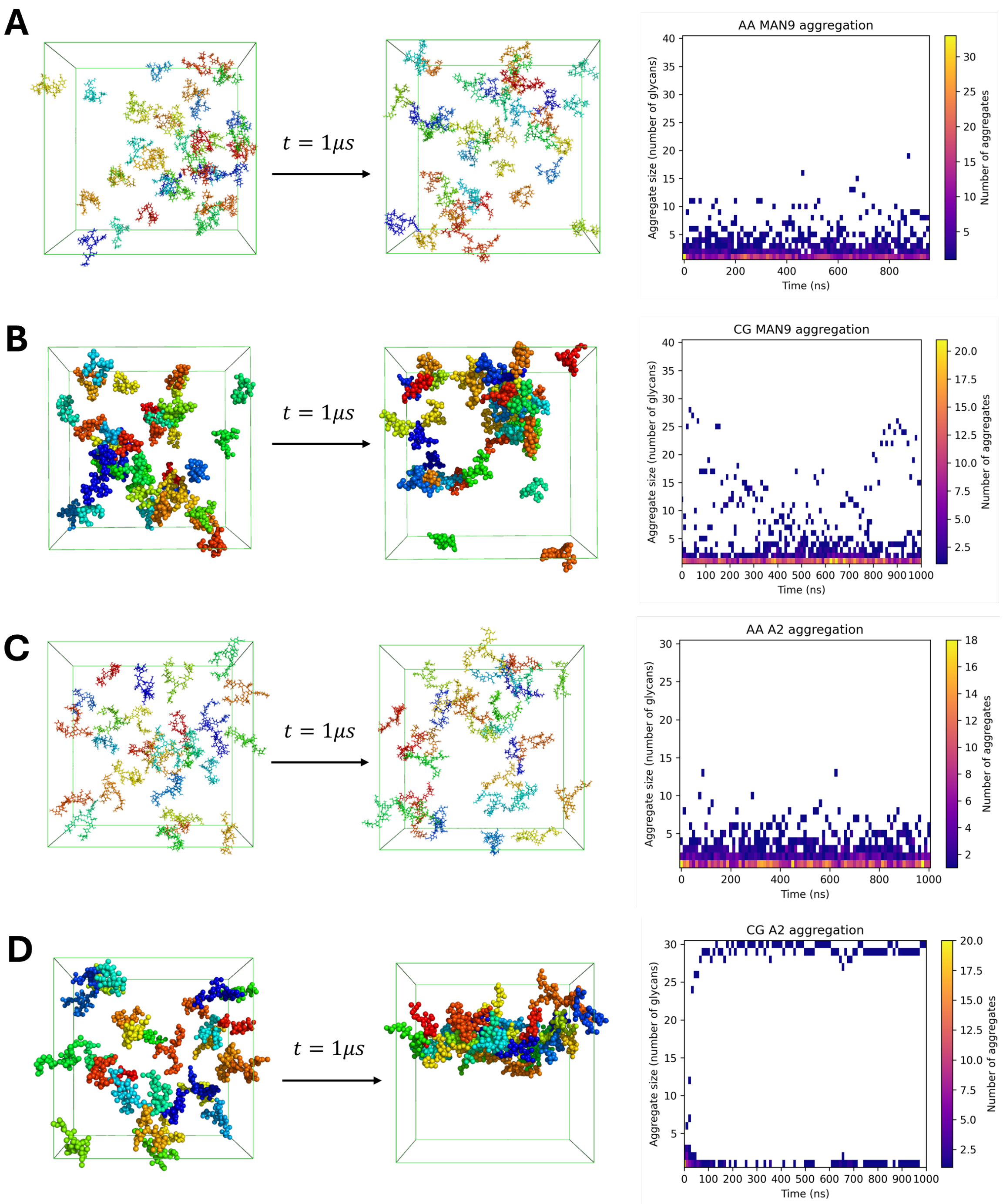


Figure S27: **Self-association of free N-glycans in solution: AA versus CG.** Forty copies of M9 (**A**, **B**) or thirty A2 molecules (**C**, **D**) were simulated for 1 $\mu$s, where each representing concentration of 50 g/L with adjusted cube box sizes, using the CHARMM36m force field (**A**, **C**) or the GMR Martini 3 models (**B**, **D**). Left and middle: initial and final configurations, with each glycan coloured individually. Right: time-resolved distribution of aggregate sizes, defined by a single-linkage cluster analysis with a 0.47 nm inter-glycan contact cutoff; the colour scale gives the number of aggregates of a given size at each time point. In the AA simulations, both glycans remain predominantly monomeric, with transient dimers and small oligomers (**A**, **C**). The CG M9 system samples larger but transient clusters that repeatedly dissolve, maintaining an equilibrium between monomers and aggregates (**B**). The CG A2 system, by contrast, collapses into a single aggregate within the first 50 ns that persists for the remainder of the simulation (**D**).
